# Tuning single-molecule ClyA nanopore tweezers for real-time tracking of the conformational dynamics of West Nile viral NS2B/NS3 protease

**DOI:** 10.1101/2024.05.14.594247

**Authors:** Spencer A. Shorkey, Yumeng Zhang, Jacqueline Sharp, Sophia Clingman, Ly Nguyen, Jianhan Chen, Min Chen

## Abstract

The flaviviral NS2B/NS3 protease is a conserved enzyme required for flavivirus replication. Its highly dynamic conformation poses major challenges but also offers opportunities for antiviral inhibition. Here, we established a nanopore tweezers-based platform to monitor NS2B/NS3 conformational dynamics in real-time. Molecular simulations coupled with electrophysiology revealed that the protease could be captured in the middle of the ClyA nanopore lumen, stabilized mainly by dynamic electrostatic interactions. We designed a new *Salmonella typhi* ClyA nanopore with enhanced nanopore/protease interaction that can resolve the open and closed states at the single-molecule level for the first time. We demonstrated that the tailored ClyA could track the conformational transitions of the West Nile NS2B/NS3 protease and unravel the conformational energy landscape of various protease constructs through population and kinetic analysis. The new ClyA-protease platform paves a way to high-throughput screening strategies for discovering new allosteric inhibitors that target the NS2B and NS3 interface.

## Introduction

Arthropod-borne flaviviral viruses, including dengue (DENV), Zika (ZIKV), West Nile (WNV), Japanese encephalitis (JEV), tick-borne encephalitis (TBEV) and Yellow fever (YFV), present an increasing threat to public health as potential epidemics due to the extensive global spread and transmission in the past decades.^1-5^ So far, there is no approved antiviral treatment against flavivirus infection, and vaccines are only available for some flaviviruses, with some vaccines only protecting limited populations in certain ages.^6^ Like other flaviviral members, the WNV genome encodes a tryp-sin-like serine protease NS2B/NS3 consisting of two core components, the NS3 protease and a NS2B co-factor peptide (Figure 1a).^7-9^ Due to its essential role in the viral life cycle^10^, NS2B/NS3 has been recognized as a high-priority drug target for developing antiviral therapies.^7, 11^ Despite extensive efforts in searching inhibitors against NS2B/NS3’s highly conserved active site, no molecule has shown sufficient drug-like properties to reach clinical trials.^12^ It is believed that the relatively shallow active site and its preference for substrates containing dibasic residues are the major obstacles. Recent efforts have shifted toward potential allosteric druggable sites at the NS2B and NS3 interaction interface.^13, 14^

**Figure 1:**
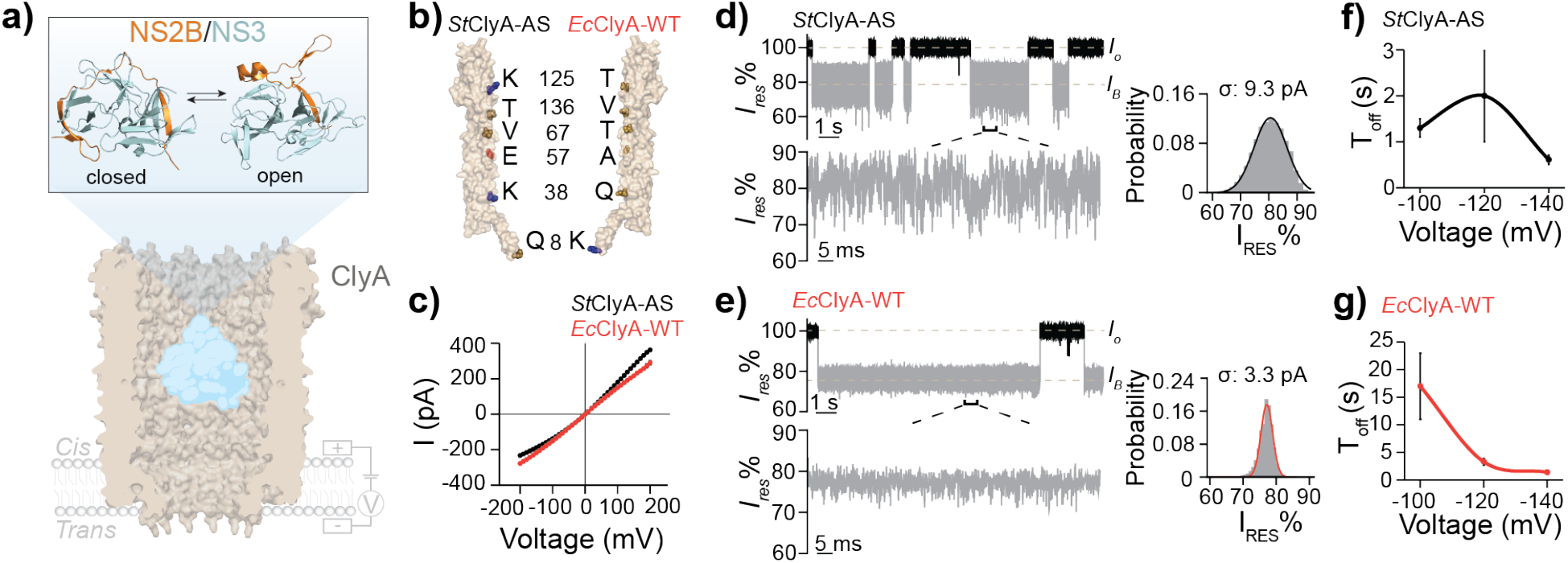
ClyA nanopores for trapping the WNV NS2B/NS3 protease. **a)** Schematic illustration of a WNV NS2B/NS3 protease trapped within ClyA nanopore tweezers (PDB: 6MRT). Structures of a closed (top left, PDB: 2FP7) and an open state (top right, PDB: 2GGV) were shown. **b)** Residues within the nanopore lumen that are different between *St*ClyA-AS (left) and *Ec*ClyA-WT (right). **c)** Current-voltage relationships of *St*ClyA-AS and *Ec*ClyA-WT nanopores. **d-e)** Representative current recording traces of the protease trapping by *St*ClyA-AS and *Ec*ClyA-WT at -120 mV. Corresponding all points histograms of all trapped events for a single trace were fitted by a Gaussian function to derive the stand deviation σ. **f-g)** Dwell times (*T*_off_) of the protease in *St*ClyA-AS and *Ec*ClyA-WT across a range of voltages.

In the ligand-free state, nuclear magnetic resonance (NMR)^15^ and small-angle X-ray scattering (SAXs)^16^ revealed significant structural dynamics within WNV NS2B/NS3, suggesting an equilibrium of multiple conformations in the apoenzyme.^11^ X-ray crystallographic studies have reported NS2B/NS3 in “closed” and “open” conformations (Figure 1).^17, 18^ In the “open” conformation, the NS2B C-terminal β hairpin (NS2BCtβ, residues 74-87) is located away from the active site, whereas the “closed” state forms a lid partially covering the catalytic region (Figure 1a).^19^ Numerous biochemical studies have demonstrated that the activity of NS2B/NS3 was strongly correlated with structural dynamics. For example, a crosslinked DENV-2 NS2B/NS3 that fixed the enzyme in a closed conformation showed a four-fold increase of catalytic activity.^20^ In contrast, mutations that destabilized interactions between NS2B and NS3 exhibited significantly reduced *k*_*cat*_.^21^ These findings suggest that locking NS2B/NS3 in the open conformation is an effective strategy to inhibit its activity.^22^ Recently, Li et al. established a cell-based assay using split-luciferase to screen for Zika NS2B/NS3 protease interface inhibitors and applied it to a moderate library comprising 2816 compounds.^23^ Nonetheless, cell-based assays are often beset by high costs, prolonged development periods, and susceptibility to off-target effects.^24^ Virtual screening for interface binders of NS2B/NS3 has been limited by a lack of reliable structural models of open states for docking tests.^25^ A cost-effective, in vitro high-throughput screening strategy capable of identifying interface binders is highly desirable to expedite the discovery of novel allosteric inhibitors.

Nanopore genomic sequencing has emerged as a powerful technology for genomics and transcriptomics due to its ability for long reads and direct detection of nucleotide modifications.^26-28^ As a label-free single-molecule technique, nanopore sensing shows significant promise across a wide spectrum of bioanalytical applications beyond DNA sequencing,^29^ particularly within the realm of protein analysis such as protein target detection,^30-34^ post-translational modification identification,^35-39^ and protein sequencing.^40-42^ Notably, nanopore tweezers have been recently developed for monitoring protein conformational changes induced by ligand-binding and catalytic activities.^43-46^ Unlike classic nanopore analysis which measures the current fluctuations induced by analytes transiently traversing the nanopore, nanopore tweezers work by confining a single analyte molecule within its lumen to continuously record ionic current fluctuations associated with the analyte’s motions in real time.^44, 47-51^ Cytolysin A (ClyA), a pore-forming protein in enterobacterial species including *Escherichia coli* (*E. coli*) and *Salmonella typhi (S. typhi)*,^52^ has been developed as a nanopore tweezers for studying the ligand binding of multiple substrate-binding proteins,^49^ resolving the conformational energy landscape of a pharmaceutically important protein, Abl kinase domain, and detecting the inhibitor imatinib binding.^44^

To select molecules which stabilize the inhibitory open conformations, it is necessary to establish an approach capable of observing the open and closed conformations in NS2B/NS3. In this work, we combined molecular dynamics (MD) simulations and single channel current measurements to guide the fine-tuning of an *S. typhi* ClyA nanopore tweezers’ resolution to achieve differentiation of the open and closed conformational transitions of WNV NS2B/NS3.

## Results

### E. coli and S. typhi ClyA exhibit distinct tweezer behaviors

Previously, an *E. coli* wild-type ClyA (*Ec*ClyA-WT) and an engineered *S. typhi* ClyA AS construct (*St*ClyA-AS), have been used as nanopore tweezers for trapping various analytes to detect protein-ligand interactions and reveal conformational changes.^44, 47, 48^ Interestingly, we discovered that *Ec*ClyA-WT could resolve three distinct maltose binding states to maltose binding protein (MBP), associated with MBP loaded with different anomeric stereoisomers of maltose.^47^ In contrast, *St*ClyA-AS did not detect multiple maltose binding states under identical experimental conditions,^53^ highlighting significant differences in sensitivity between the two nanopores with 88% sequence identity (Supplemental Figure S1). Distinct residues located at the pore lumen of the two nanopores are indicated in Figure 1b. For comparison, *St*ClyA-AS has a slightly more hydrophilic/charged lumen than *Ec*ClyA-WT, while the net charges of the two pores’ lumen are identical. Both ClyA nanopores exhibited a broad distribution of open pore conductance due to their ability to form multimers of different stoichiometry.^54, 55^ Nevertheless, the current-voltage curves of two pores of the same conductance (as determined at 20 mV) are very similar at low voltages (Figure 1c), while at high voltages, *Ec*ClyA-WT has a more linear I-V relationship than *St*ClyA-AS, which exhibits rectification.

Both *Ec*ClyA-WT and *St*ClyA-AS pores can trap the West Nile NS2B/NS3 viral protease but show different current signals and trapping lifetimes. Specifically, NS2B/NS3 in *St*ClyA-AS exhibits dynamic current blockades with 70-90% residue current (*I*_res_, defined as the current of blockades *I*_*B*_ divided by the open pore current *I*_*o*_) with a standard deviation σ of 9.6 pA, whereas NS2B/NS3 in *Ec*ClyA-WT shows relatively quieter blockades from 70-80% *I*_res_ with an σ of 3.1 pA (Figure 1d, e, Supplemental Figure S2). At -100 mV, NS2B/NS3 exhibits more than ten times longer dwell times in *Ec*ClyA-WT relative to *St*ClyA-AS nanopore (Fig.1f, g). Furthermore, the dwell time first increases up to -120 mV and then decreases at -140 mV in *St*ClyA-AS, whereas for *Ec*ClyA-WT, the dwell time maxes at -100 mV and decreases significantly at greater voltages. These results indicate that the NS2B/NS3 protease can translocate through both ClyA nanopores at high voltages. When trapped in ClyA, NS2B/NS3 is controlled by three forces, including electrophoresis, electroosmosis, and prote-ase-nanopore interactions. The theoretical isoelectric point (pI) of NS2B/NS3 is 5.2. Under the experimental conditions, electroosmosis drives NS2B/NS3 into the pore, against electrophoresis, balancing with the protease-nanopore interactions to stabilize the protease at a specific location within the lumen. Due to high sequence identity, the two nanopores are expected to have essentially the same structures and almost identical net surface charge in the lumen. Thus, the significant difference in dwell time and current trace fluctuations observed between *St*ClyA-AS and *Ec*ClyA-WT is surprising. We postulate that the divergent current signals may be due to differences in the interaction between the protease and the nanopore.

### Molecular modeling reveals a key mode of dynamic protease-ClyA interactions

To probe for the molecular basis on why *St*ClyA-AS and *Ec*ClyA-WT might be different in their interactions with the NS2B/NS3 protease, we first conducted steered molecular dynamics (MD) simulations where the protease was gradually pulled into the ClyA pore (Supplemental Figure S3). The system was described using a hybrid resolution (HyRes) coarse-grained protein model^56, 57^, which has been previously shown to be able to capture dynamic protein interactions and phase transitions.^58,59^ Based on prior estimates,^60^ the pulling force in the simulation was set to 60 pN, which is large enough to drive the protease to the end of the barrel within 50 ns in most cases but also small enough to probe potential interactions of the protease with the inner wall of the ClyA lumen. Interestingly, as NS2B/NS3 was pulled through the ClyA pore toward the constriction region, we frequently observed dynamic interactions between the protease and the ClyA pore lumen residues (Figure 2a, Supplemental Figure S4). This observation indicated the existence of stable interactions formed between the protease and ClyA about halfway down the nanopore barrel (around K147), leading to “mid-trap” states that could persist throughout the simulation time span (e.g., Figure 2a bottom panel). The mid-trap states existed for both *St*ClyA-AS and *Ec*ClyA-WT, regardless of whether the protease was simulated in the open or closed state.

**Figure 2:**
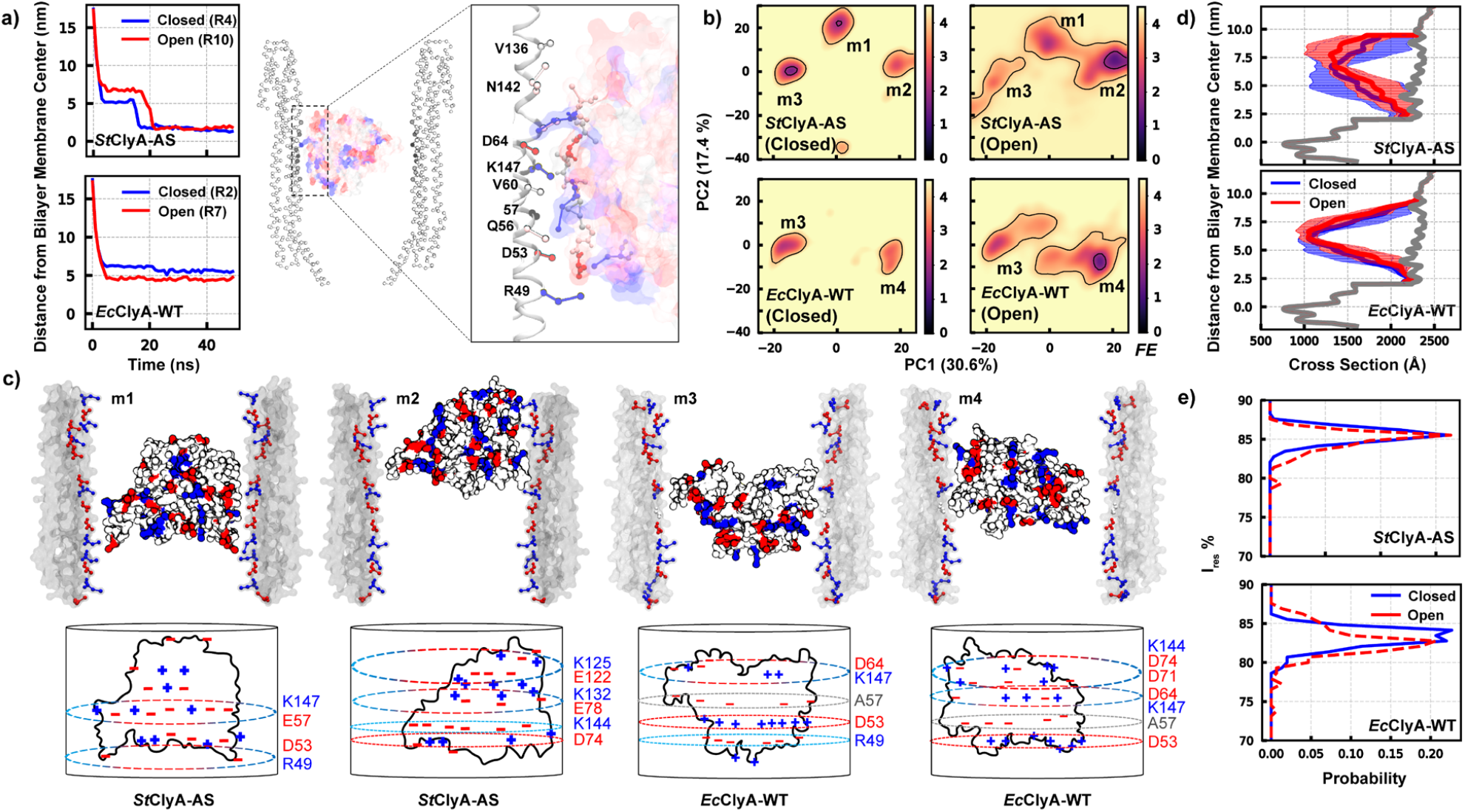
Dynamic interactions of the WNV NS2B/NS3 protease with *S. typhi* ClyA mutant pores. **a)** Distributions of the protease conformational ensembles in the mutant ClyA nanopore in the mid-trap state, projected along the same first two principal components as used in Figure 2b. See Figure 2b caption for the plotting scheme. **b)** Averaged pore opening profiles of *St*ClyA-E57A (top) and *St*ClyA-E57K (down) pores with an Open (red) and Closed (blue) protease captured in the mid-trap state. The shaded areas indicate the standard deviations derived from MD ensembles. The gray traces correspond to the profiles of the empty pore. For protease dynamics in StClyA-E57A pore, only trajectories contributing to m2 binding states were used to conduct the cross-section and *I*_res_% calculations. **c)** The probability distributions of *I*_*res*_% of Open (red) and Closed (blue) protease in *St*ClyA-E57A (top) and *St*ClyA-E57K (down) pores estimated from the cross-section areas (see Methods).

To further characterize the nature of the mid-trap states, we preformed additional MD simulations in the absence of the pulling force, with the protease initially placed in the middle of the nanopore in random orientations. The distributions of resulting ensembles are summarized in Figure 2b, with key states illustrated in Figure 2c and Supplemental Figure S5. The results reveal substantial heterogeneity in the configuration of the protease-ClyA interactions and the existence of multiple free energy basins in all four cases. The protease apparently samples a broader configurational space in the open state, which is likely a direct consequence of larger conformational flexibility of the protease itself. Importantly, the protease, regardless of whether it is in the open or closed states, appears more dynamic and samples more sub-states inside the *St*ClyA-AS pore than it does inside *Ec*ClyA-WT. This observation is consistent with the larger standard deviation of current blockage (∼9.6 pA in *St*ClyA-AS vs. ∼3.1 pA in *Ec*ClyA-WT, Figure 1 d, e). The *I*_res_% directly estimated from the pore opening cross-section profiles (Figure 2d) suggest that the mid-trap states would give rise to 80-85% blockage in both pores (Figure 2e), which is in general agreement with the current measurements (Figure 1). We note that blockage percentages estimated directly from cross-section areas may not be quantitatively accurate.^61^

Further analysis revealed that electrostatic interactions play a major role in stabilizing the protease/ClyA interaction (Figure 2c). The interior wall of the ClyA nanopore contains several rings of charged residues, such as D63, K147, D53 and R49. Interestingly, multiple orientations exist where the protease can present charged belts on its surface to match the charge rings on ClyA, which are exemplified the best by the m1 and m3 substates (Figure 2c). Examination of the charge complementarity in these states also reveals how a single sequence difference in this region of the ClyA interior pore surface (E57 in *St*ClyA-AS vs A57 in *Ec*ClyA-WT) could result in divergent current signals measured. The presence of an extra negative charge ring of E57 in *St*ClyA-AS would lead to unfavorable repulsion in substates m3 (as well as m4 observed in *Ec*ClyA-WT), where a belt of negative charges on the protease is positioned to interact with the residue 57 region. As a result, the protease is apparently pushed higher and samples additional states including m1 and m2. With a neutral side chain of A57 in *Ec*ClyA-WT, both substates m3 and m4 are much more stable and dominate the dynamic interactions, leading to quieter blockade signals.

### Design of tailored ClyA nanopores for monitoring NS2B/NS3 conformational dynamics

To probe the importance of position 57 in the predicted mid-trap states, we designed two *St*ClyA-AS mutant pores, *St*ClyA-E57A and *St*ClyA-E57K, and simulated their dynamic interactions of the NS2B/NS3 protease. The results, summarized in Figure 3, suggest that both E57A and E57K mutations lead to more orderly binding configurations in the mid-trap states. Neutralization of E57 with alanine removes the negative charge repulsion and both m3 and m4 become much more stable, even though substantial dynamics appear to persist in the open and more flexible state of the protease (Figure 3a). With *St*ClyA-E57K, the m3 substate becomes the most dominant one for both the open and closed states of the protease. Correspondingly, *I*_res_% estimated from the open cross-section profiles (Figure 3b) suggest that the *St*ClyA-E57K pore will not likely be able to resolve the open and closed states (Figure 3c). Instead, the levels of blockade of the open and closed states of the protease may be more discernable in the *St*ClyA-E57A pore, with the open state estimated to yield ∼3% lower *I*_res_.

**Figure 3:**
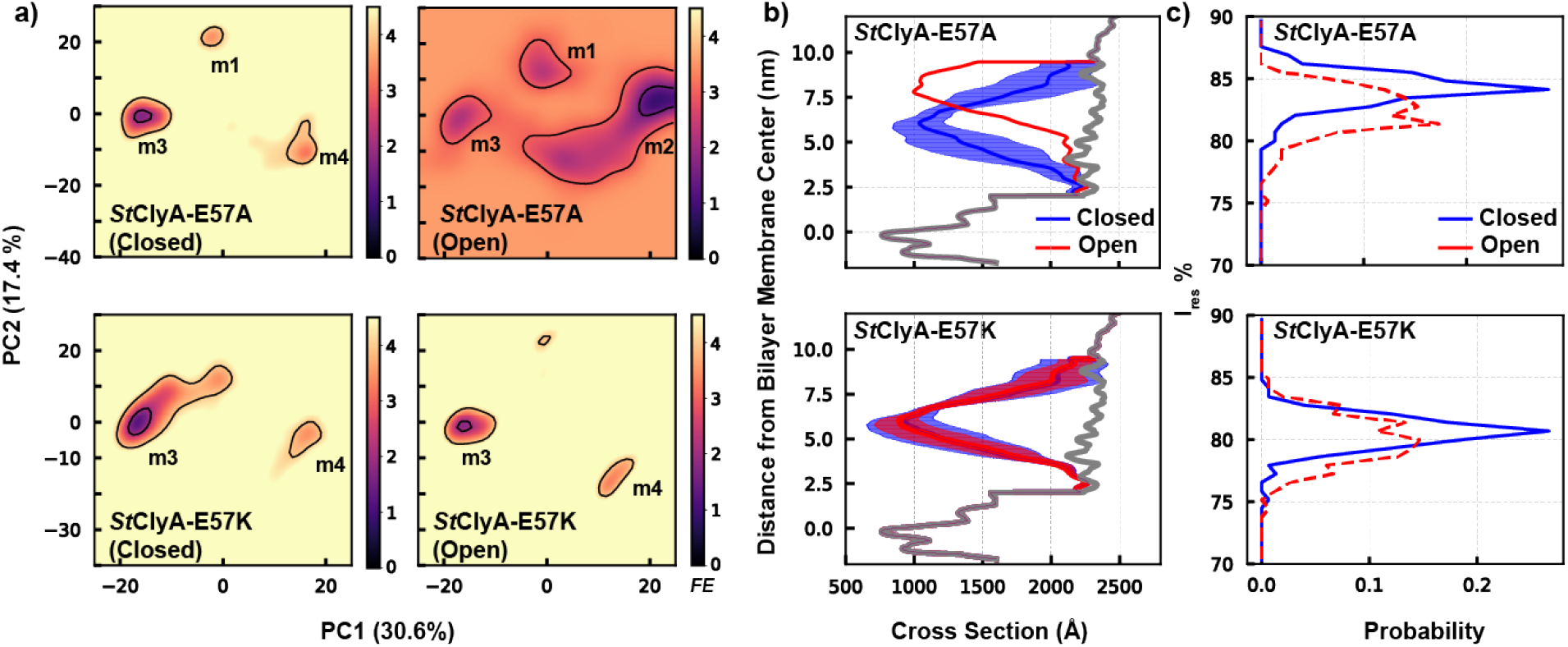
Dynamic interactions of the WNV NS2B/NS3 protease with S. *typhi* ClyA mutant pores. **a)** Distributions of the protease conformational ensembles in the mutant ClyA nanopore in the mid-trap state, projected along the same first two principal components as used in Figure 2b. See Figure 2b caption for the plotting scheme. **b)** Averaged pore opening profiles of *St*ClyA-E57A (top) and *St*ClyA-E57K (down) pores with an Open (red) and Closed (blue) protease captured in the mid-trap state. The shaded areas indicate the standard deviations derived from MD ensembles. The gray traces correspond to the profiles of the empty pore. For protease dynamics in StClyA-E57A pore, only trajectories contributing to m2 binding states were used to conduct the cross-section and *I*res% calculations. **c)** The probability distributions of *I*res% of Open (red) and Closed (blue) protease in StClyA-E57A (top) and StClyA-E57K (down) pores estimated from the cross-section areas (see Methods).

To test the predicted mutational effects, we generated two nanopores, *St*ClyA-E57A and *St*ClyA-E57K, and assessed their ability to trap the protease. Both mutant pores successfully trapped NS2B/NS3 showing 80% *I*_res_, with noise levels reduced threefold compared to *St*ClyA-AS (Figure 4a). Remarkably, the current noise σ values for the two *St*ClyA-AS mutants closely resemble that of *Ec-* ClyA-WT, with *St*ClyA-E57K displaying the smallest σ of 2.9 ± 0.1 pA. This result strongly supported the observation from MD simulations that residue 57 of ClyA was directly involved in interacting with the trapped protease, because mutations at this site significantly altered the noise levels of current signals as predicted. Our current recording data and MD simulation synergistically support that the negatively charged glutamate E57 at *St*ClyA-AS destabilizes the protease-nanopore interaction via charge repulsion, leading to greater conformational heterogeneity and higher current noise. Interestingly, E57A and E57K mutations exhibited opposite effects on the dwell time of the protease (Figure 4b). *St*ClyA-E57A slightly increased the dwell time, whereas *St*ClyA-E57K significantly reduced it. Both mutations were anticipated to weaken electroosmosis, leading to reduced dwell times. The interevent time τ_on_ analysis showed that *St*ClyA-E57K increases τ_on_ approximately four-fold, while *St*ClyA-E57A has a τ_on_ similar to that of *St*ClyA-AS (Figure 4c). Given that the τ_on_ of the trapping events is mainly governed by the electroosmotic flow, these results strongly support that *St*ClyA-E57K significantly diminishes the electroosmotic force, resulting in a short dwell time of NS2B/NS3 in this nanopore.

**Figure 4:**
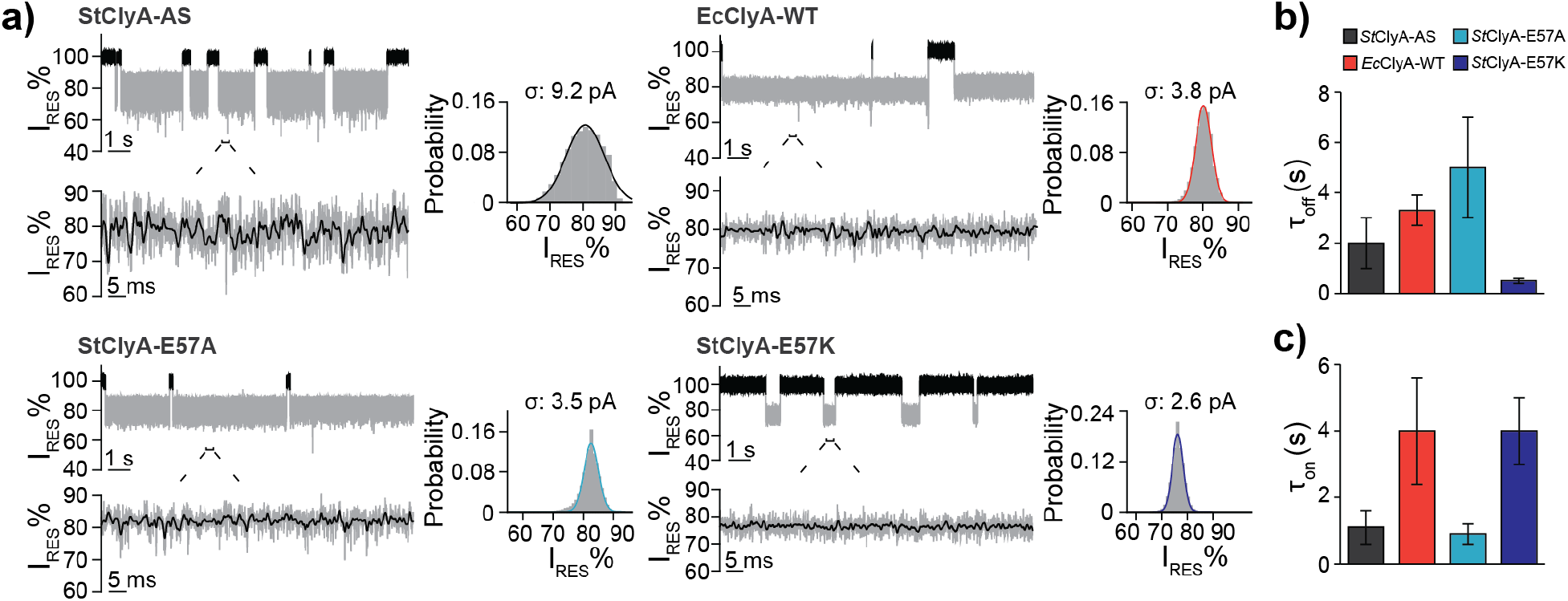
Characterization of mutant *S. typhi* ClyA pores. **a)** Representative current traces and the corresponding all-points histograms of NS2B/NS3 protease trapped by *St*ClyA-AS, *Ec*ClyA-WT, *St*ClyA-E57A, and *St*ClyA-E57K at -120 mV. **b)** Dwell times (τ_off_) and **c)** Inter-event times (τ_on_) of the protease at 200 nM in each ClyA pore variant at -120 mV. The error bars were derived from analysis of at least three separate pores.

In summary, our computational and experimental findings confirm the central role of the midpore region around residue 57 of ClyA in interacting with NS2B/NS3. Despite both *Ec*ClyA-WT and *St*ClyA-E57A nanopore constructs exhibiting similar low noise levels, *Ec*ClyA-WT has a lower voltage tolerance than *St*ClyA, as *Ec*ClyA-WT typically starts to gate frequently at voltages higher than -80 mV whereas *St*ClyA can stay open mostly at -120 mV. Thus, the *St*ClyA-E57A construct exhibits optimal nanopore tweezer properties, characterized by prolonged NS2B/NS3 dwell times, low signal noises, a fast capture rate and high voltage tolerance.

### Probing the open and closed conformational dynamics of the NS2B/NS3 protease

We investigated using *St*ClyA-E57A to probe the open and closed conformational dynamics of NS2B/NS3. To correlate observed current levels with structural states, we generated four NS2B/NS3 mutants to perturb the open and closed conformational equilibrium, including (Figure 5a): 1) NS3 C78A, referred to as wild type (WT) in this section, which removes the single cysteine while remaining active (Supplementary Figure S6). 2) NS3-T111F, denoted as T111F, which disrupts a hydrogen bond between NS3-T111 and NS2B-T69 and results in a protease with significantly decreased activity (Supplementary Figure S6).^21^ 3) Double cysteine mutant NS2B-D76C*/NS3-K117C-C78A in the reduced form, referred to as rd-C76*/C117 (* denotes residues on NS2B), disrupting a key salt-bridge formed in the closed conformation between NS2B-D76* and NS3-K117. 4) NS2B-C76*/NS3-C117-C78A protein with two cysteines crosslinked to lock the protease in a closed conformation, termed as xc-C76*/C117.

**Figure 5:**
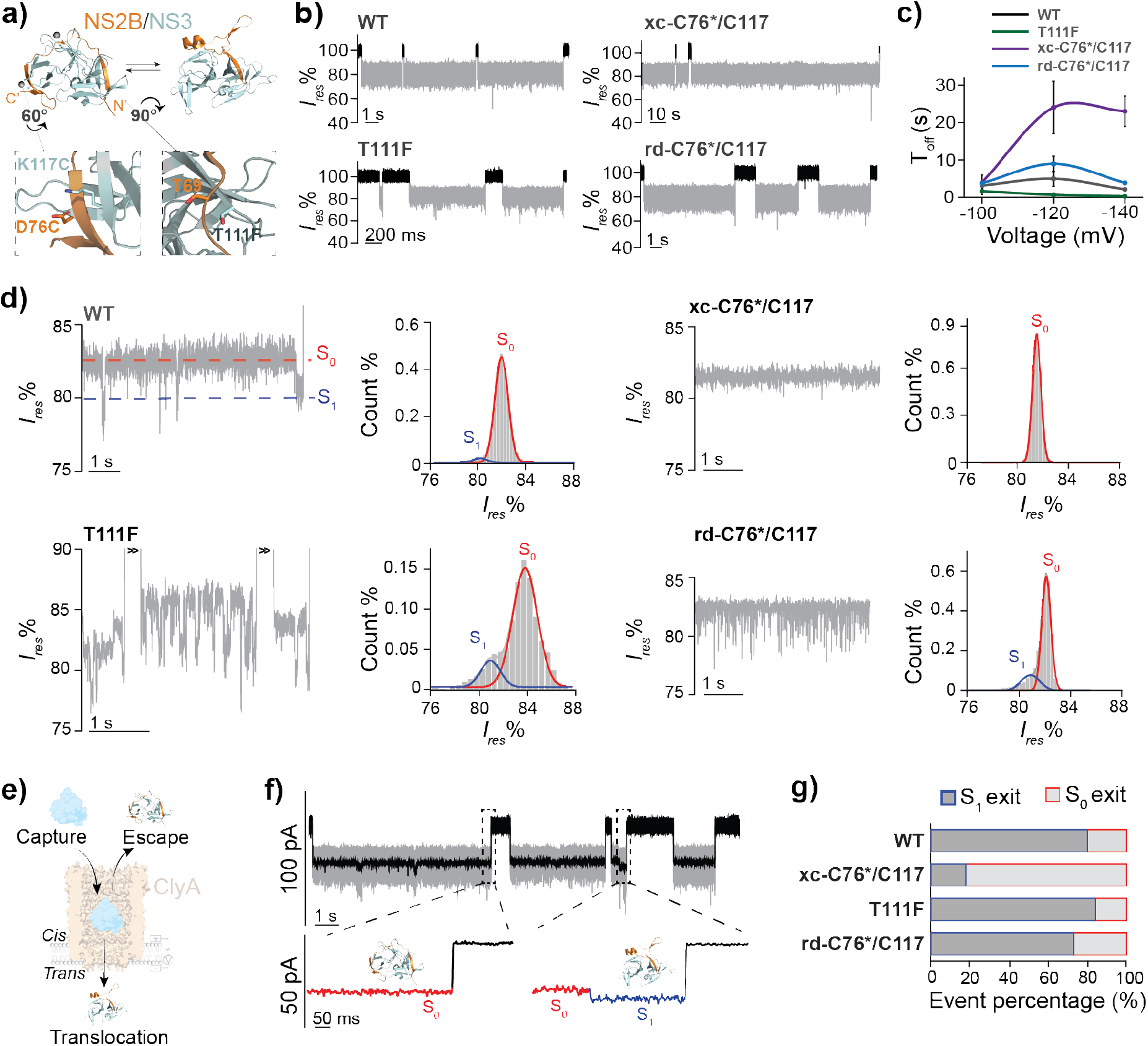
Detection of the open and closed conformational states of the WNV NS2B/NS3 protease. **a)** Structures of WNV NS2B/NS3 showing the closed (top left, PDB: 2FP7) and open states (top right, PDB: 2GGV). Mutation sites D76C* of NS2B, K117C and T111F of NS3 were labelled in the bottom panels. **b)** Representative current traces of each NS2B/NS3 construct trapping in the *St*ClyA-E57A pore at -120 mV. **c)** Dwell times (*T*_off_) of NS2B/NS3 variants at various voltages. **d)** Current traces of NS2B/NS3 variants filtered by a 100 Hz low-pass Gaussian filter. Traces of three individual trapped events of T111F were shown due to their short trapping times. Corresponding histograms were fitted with Gaussian functions to derive the populations of S_0_ and S_1_ states. The standard deviation σ of S0 noises of filtered traces is 0.77 ± 0.12, 0.97 ± 0.09, 0.56 ± 0.10, and 0.55 ± 0.11 pA for WT, T111F, xc-C76*/C117, and rd-C76*/C117, respectively. As a reference, the σ for the unoccupied ClyA pore was 0.49 ± 0.11 pA. **e)** Schematic of the NS2B/NS3 protease entering and exit from ClyA (PDB: 6MRT). **f)** A representative current recording trace of WT NS2B/NS3 showcasing two different exit modes. **g)** Fractions of NS2B/NS3 variants existing in S_0_ or S_1_ states. For each construct, the exit states were assigned for all events of three pores and summed to determine the total number of events for which the protease occupied either the S_0_ or S_1_ state prior to exiting the pore. All current measurements were performed in 20 mM HEPES pH 7.4 150 mM NaCl. Data were acquired at -120 mV with a 50 kHz Bessel filter at a sample rate of 5 kHz.

All four proteases trapped by *St*ClyA-E57A exhibit current blockades with an average *I*_res_ of approximately 82% (Figure 5b, Supplementary Figure S7). Interestingly, they show different dwell times despite having identical net charges (Figure 5c). The trapping times of rd-C76*/C117 and WT slightly increase from -100 mV to -120 mV, then decrease at -140 mV, while T111F shows a decreased trapping time over -100 mV to -140 mV. This observation indicates the translocation of these proteins to the trans side of the pore at high voltages. In contrast, xc-C76*/C117 exhibits a substantially increased dwell time compared to other pores, and show no obvious reduction at -140 mV, suggesting it cannot pass through the nanopore.

After filtering the traces with a 100 Hz low-pass Gaussian filter, two current states with a ∼2-6 pA difference were observed for WT (Figure 5d). The higher residual current state, S_0_, is the dominant one with a 92.6 ± 9.1 % population occupancy. In contrast, the xc-C76*/C117 construct completely abolishes S_1_ states, suggesting that S_1_ states in WT are associated with open conformations (Figure 5d). This assignment is in agreement with the prediction from MD simulations, that *I*_res_ for the open state in *St*ClyA-E57A should be smaller (Figure 3c). Indeed, the destabilization mutant T111F exhibits more frequent S_0_/S_1_ transitions and reduces S_0_ occupancy to 81.3±3.4% compared to WT, which is consistent with reduced activity of T111F.^21^ Mutant rd-C76*/C117 also shows more frequent transitions between S_0_ and S_1_ states, with the S_0_ occupancy decreased to 79.1±8.2%, which is consistent with the disruption of a key closed state salt-bridge in this construct. Together, these observations strongly support that the S_0_ and S_1_ current states in *St*ClyA-E57A correlate with the closed and open conformational states of NS2B/NS3, respectively.

Once a protein is captured by ClyA, it can either escape to the cis side or translocate through the pore to the trans side (Figure 5e). We observed two distinct types of current signals, S_0_ or S_1_, at the end of each trapping event (Figure 5f). Strikingly, ∼80% of events in WT, T111F, and rd-C76*/C117 have S_1_-type endings, in stark contrast to the ∼10% for the crosslinked protease xc-C76*/C117 (Figure 5g). Since xc-C76*/C117 cannot translocate through the pore, we interpreted this result as the S_1_-ending events being concluded by translocation, while S_0_-ending events are likely the “escape” events. This explanation is supported by the significantly longer dwell time of xc-C76*/C117 (Figure 5c) than those of other three protease constructs. From a structural point of view, the rigid closed conformation of the crosslinked protein prohibits translocation. Conversely, WT, T111F, and rd-C76*/C117 could switch to more flexible S_1_ open states, facilitating the enzyme squeezing through the constriction site of the pore. Thus, the different preferences of these protease constructs to exit the pore provide additional evidence to support that the S_1_ states are associated with open conformations of the protein complex and S_0_ the closed one.

In summary, our *St*ClyA-E57A pore can detect and differentiate between the protease’s open and closed conformations manifested as the S_1_ and S_0_ current states in current recordings.

## Discussion

### The energy landscape of NS2B/NS3 structural dynamics

Previous NMR studies have revealed that the major population (>90%) of NS2B adopts a closed state, along with minor contributions (∼5%) of relatively disordered “open” states, where NS2B could adopt multiple positions relative to NS3.^15^ SAXS studies align with this observation, suggesting 85% of the NS2B/NS3 in a compact conformation and ∼15% in a more extended structure.^16^ Our nanopore measurement supports a similar equilibrium of predominantly closed and occasional open structural states, with the ∼92.6 % closed state population in quantitative agreement with both NMR and SAXS results. More importantly, real-time nanopore measurements provided insights into the kinetics of the open-closed transitions. The residence time distribution of S_0_ for WT exhibited two populations, a fast transition at ∼3 ms scale and a prolonged transition at 100-400 ms scale (Figure 6a). Interestingly, despite their similar closed state S_0_ occupancies, the kinetics of the open and closed transition of the two mutants, T111F and rd-C76*/C117, were drastically different. For example, the S_0_/S_1_ transitions in T111F had one rapid and one slow process suggesting two open conformations with distinct stability. In contrast, rd-C76*/C117 only showed a single fast S_0_/S_1_ transition process (Figure 6b), with a dwell time of 32.2 ± 9.9 ms and 3.0 ± 0.6 ms for S_0_ and S_1_ respectively (N=3, n=1017).

**Figure 6.**
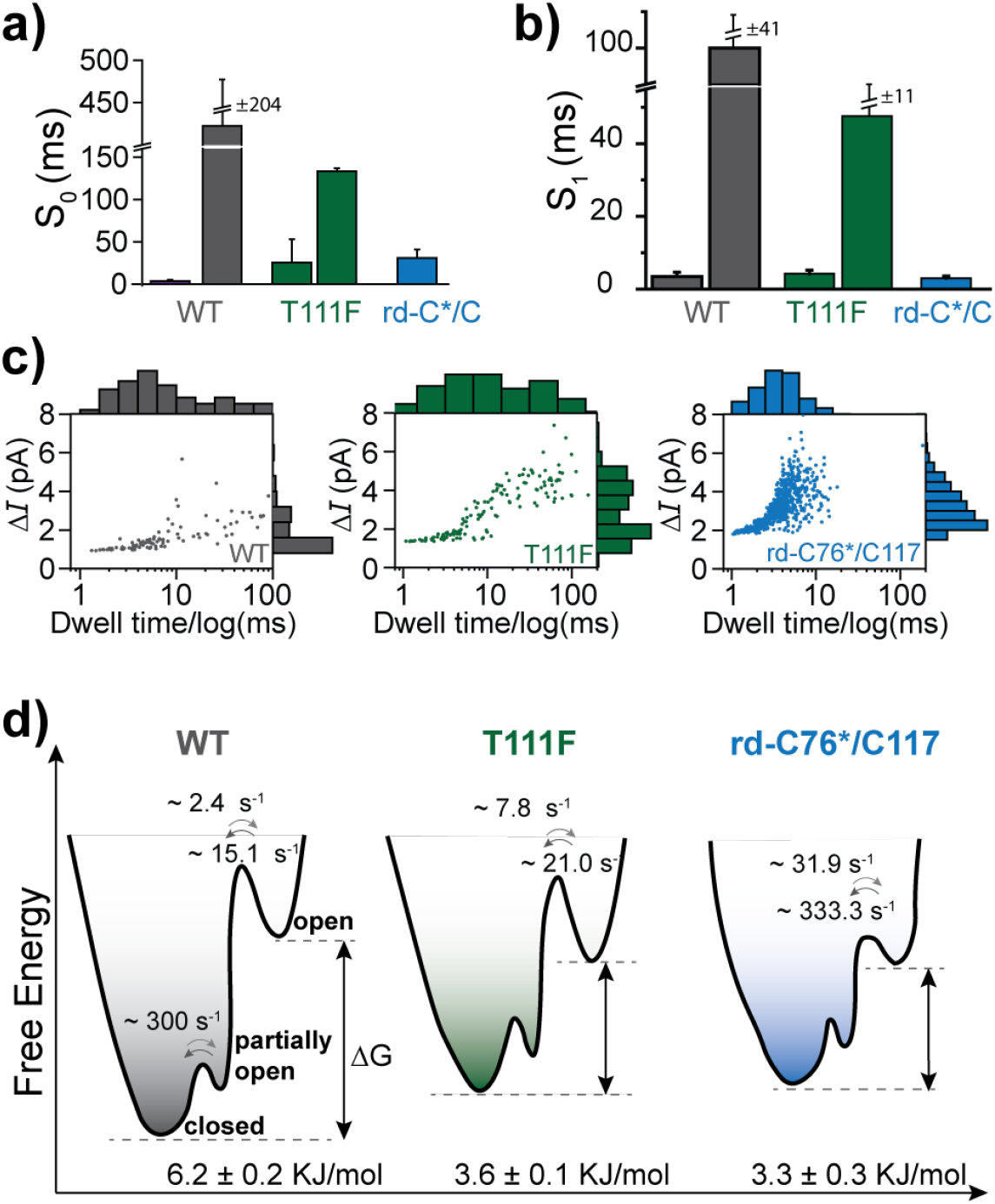
The energy landscape of WNV NS2B/NS3 proteases. **a-b**) Dwell times of S_0_ and S_1_ states for each NS2B/NS3 construct. Dwell times of WT and T111F from each pore were fitted with double exponential functions and those of rd-C76*/C117 with single exponential function (Supplemental Figure S12). **c)** 2-D event distributions of S_1_ events. **d)** The energy landscape of open and closed conformational states. Interconversion rates were estimated from the dwell times of the S_0_ and S_1_ states of each protease variants.

Two-dimensional distribution plots of S_1_ events revealed a correlation between event durations and current amplitudes *ΔI* (|*I*_s0_-*I*_s1_|) (Figure 6c). Specifically, events with shorter dwell time had a lower *ΔI*, while long-lasting events demonstrated higher *ΔI*. Analysis of the standard deviation σ of the S_0_ current signals show the T111F had an obviously increased σ compared with that of WT, while the crosslinked mutant xc-C76*/C117 strongly suppressed σ (Figure 5d), suggesting that S_0_ noises also reflect the protease’s opening motions. Collectively, these data suggest that high-frequency S_0_ noises and transient S_1_ events with small amplitudes originate from rapidly detaching only a few NS2 C-terminal amino acids and swiftly returning to the closed state, presumably due to the inability to overcome a high energy barrier created by the D76*/K117 salt bridge.

Based on S_0_/S_1_ population and kinetic analyses, we propose a conformational energy landscape for the NS2B/NS3 protease and its mutants (Figure 6d). The WT NS2B/NS3 exhibits two types of opening motions: a small amplitude, partially opening without breaking the D76*/K117 salt bridge lasting less than 10 ms, and larger conformational changes leading to open states at the 10-500 ms scale. Both T111F and rd-C76*/C117 alter the landscape to favor a more open state; however, T111F primarily destabilized the closed state, with a minimal effect on the stability of the open state. Conversely, rd-C76*/C117 significantly destabilized both the open and closed states, with the latter to a greater extent. The faster closed-to-open transition rate observed in rd-C76*/C117 compared to T111F can explained as follows: to switch to the open state, the NS2B C-terminal β-hairpin needs to detach from NS3, which requires breaking the D76*/K117 salt-bridge. C76*/C117 mutations directly decrease the energy barrier that still exists in T111F leading to a slower on-rate. Although a vast majority of S_1_ events in rd-C76*/C117 were short-lived, the real-time single-molecule nanopore measurement allowed us to observe several rare (∼3/1000 events) long-dwell time S_1_ events, demonstrating that the mutant can form a stable open conformation (Supplementary Figure S8). Nevertheless, it is unclear why rd-C76*/C117 considerably destabilized the open states. One possibility is that the residue NS2B D76 may be involved in interactions necessary for stable open conformations.

### MD-aided fine turning enzyme-nanopore interaction interface

Our results underscore the significance of the analyte protein-nanopore interaction interface in determining the nanopore’s capability to reveal subtle structural differences of the analyte protein. Once an analyte enters the ClyA lumen, it reaches an “electrokinetic” energy minimum state, where the analyte’s position and orientation are governed collectively by electroosmosis, electrophoresis, steric hindrance, and analyte-nanopore interactions. When the analyte’s conformational state is altered, the electrokinetic energy landscape may shift, changing the positioning or orientation of the analyte in ClyA. Since the degree of current blocked by the analyte in the nanopore is related to the analyte’s position, each distinct electrokinetic energy minimum could exhibit a unique current blockade level, allowing the differentiation of the protein conformational states. However, there may be more than one electrokinetic energy minimum accessible to the analyte protein trapped in the ClyA lumen. In this case, a single conformational state of the analyte could induce current signals that either oscillate between two or more discernable current levels at low frequency (∼1 Hz) as previously shown for a glutamine substrate binding protein trapped in *St*ClyA-AS,^62^ or exhibit unresolvable high frequency (∼100-1000 Hz) noises present in this study for NS2B/NS3 in *St*ClyA-AS. The kinetics of the transition of the protein between the energy minimum states are dependent on the relative stability of the analyte protein-nanopore interactions within the collective electrokinetic energy landscape.

When using nanopore tweezers to track the conformational changes of analyte proteins, such multi-energy minima present an undesirable hurdle preventing reliable differentiation of the protein conformational states. Our study here demonstrates a feasible solution of using MD simulations to guide the fine tuning of analyte-nanopore interactions to overcome the challenge. Atomistic simulations in explicit solvent have generally been required to reliably probe transient and dynamic interactions of the protein with the nanopore,^63, 64^ which, however, suffers from expensive computational costs and poor convergence in resolving dynamic analyte-nanopore interaction modes. In this work, we utilized the highly efficient HyRes coarse-grained protein force field,^56, 57^ which has been successfully used to describe dynamic protein-protein interactions and protein phase separation.^58, 59^ The success demonstrated in this work supports the promise of HyRes as a viable tool for guiding nanopore engineering.

Our results have thus clarified a fundamental principle of applying nanopore tweezers to study dynamic protein structures: the sensitivity and resolution of nanopore tweezers for discerning conformational changes are highly dependent on the positioning of the analyte and how it interacts with the nanopore lumen. While previous studies of MBP with *Ec*ClyA-WT and *St*ClyA-AS revealed the different outcomes of the two pores,^47, 53^ our work here demonstrated that even a single amino acid substitution on the ClyA lumen can substantially affect its performance. Therefore, we propose that it is possible to tune the sensitivity of nanopore tweezers to reveal subtle structural differences among conformers through multiple strategies, such as altering the chemical properties of the pore, reorienting the analyte by adjusting its dipole moment, or fitting the analyte to pores with different geometry and shapes.

## Conclusion

Combining MD simulations and electrophysiology measurements, we have resolved the molecular basis of trapping the WNV NS2B/NS3 protease in ClyA nanopores, which guided us to fine-tune the enzyme-ClyA interaction interface, enabling the clear differentiation of the open and closed states of the NS2B/NS3 protease. The resulting *St*ClyA-E57A nanopore tweezers platform allows us to monitor protease conformational transitions in real-time at the single-molecule level, revealing the energy landscape of the structural dynamics of various WNV NS2B/NS3 constructs. The methodology and principles established here will guide the design of new nanopores for resolving subtle or heterogeneous structural dynamics of biomolecules that have proven challenging for conventional techniques. Furthermore, the *St*ClyA-E57A nanopore platform developed here presents new opportunities to develop an *in vitro* high-throughput approach currently not available for identifying novel inhibitory molecules that can stabilize the open conformational state to suppress protease activity, such as by intercalating into the NS2B and NS3 interface.

## Author Contributions

S.A.S. designed, performed biochemical and biophysical experiments, and wrote the manuscript. Y.Z. performed the MD simulation and wrote the manuscript. J.S. performed the data analysis and prepared the figures. J.S. and S.C assisted in current recording experiments. J.S. and L.N. assisted in the biochemical experiments. J.C. and M.C. wrote and edited the manuscript and supervised the project.

## Acknowledgments

This research was supported by the US National Institutes of Health grants R01AI156187 (to M.C.) and R35 GM144045 (to J.C.).

## Materials and Methods Materials

All chemicals were purchased from Thermo Fisher Scientific unless otherwise specified. 2XYT medium, DTT, and LB agar were obtained from Boston BioProducts. Pyr-RTKR-AMC was acquired from Bachem. Ampicillin and acrylamide/bis-acrylamide (29:1 ratio solution) were purchased from Research Products International. Kanamycin was obtained from Sigma-Aldrich. Agarose LE and DDM were purchased from Goldbio. Plasmid purification miniprep kits were acquired from New England Biolabs. Mini-PROTEAN TGX Stain-Free gels were purchased from BioRad. BL21(DE3) pLysS chemically competent *E. coli* cells were acquired from Invitrogen. 1,2-Diphytanoyl-sn-glycerol-3-phos-phocholine (DPhPC) was obtained from Avanti Polar Lipids, USA. Oligos were purchased from Eurofins.

### Cloning of plasmid constructs containing protein ClyA, NS2B/NS3, and TEV protease

The *E. coli* ClyA gene was amplified from the *E. coli* K-12 genome (ATCC) and cloned into a pT7 vector containing an ampicillin resistance gene and a C-terminal hexa-histidine tag as previously described.^54^ The *S. typhi* ClyA-AS sequence, as previously described,^49^ was synthesized and cloned into the pET28b vector containing an ampicillin resistance gene by Gene Universal (Newark, DE). Protein coding sequences for *E. coli* and *S. typhi* AS ClyA are shown in Supplemental Figure 1. The StAS-E57A and StAS-E57K mutants were generated using overlapping PCR with the primers listed in Supplemental Table 1.

The WNV NS2B/NS3 gene in a pET28 vector containing kanamycin resistance gene was received as a generous gift from Prof. Radhakrishnan Padmanabhan at Georgetown University. The construct was modified using overlapping PCR to include an N-terminal hexa-histidine tag (his-tag) followed by a TEV cleavage site at NS2B, and a G4SG4 linker was added to connect the C-terminus of NS2B to the N-terminus of NS3, replacing the KR dibasic auto-cleavage site. Finally, the C78A mutation was introduced to NS3 to create a cysteine-less NS2B/NS3 construct, referred to as the wild-type protease (WT) in our study. Two other mutants, NS2B/NS3-T111F and NS2B-D76C*/NS3-K117C constructs were generated by overlapping mutagenesis PCR using the cysteine-less WT construct as the template. The sequences of primers are indicated in Supplemental Table 1. The sequences of all protease variants are listed in Supplemental Table 2.

A truncated form of the Tobacco Etch Virus (TEV) protease gene was obtained as a gift from Prof. Eric Streiter as previously described.^65^

### Expression and purification of ClyA, NS2B/NS3 and TEV

Plasmids containing ClyA, NS2B/NS3, or TEV protease constructs were transformed into BL21(DE3) competent *E. coli* cells. Cells from a single colony were initially grown in 15 mL 2XYT media under ampicillin (ClyA or TEV protease) or kanamycin (NS2B/NS3) selection, shaking at 200 rpm, at 37 °C. After 6-8 hours of growth, this starter culture was transferred to 0.5 L 2XYT with 5 g/L glycerol and 2 g/L lactose and incubated for 16 hours at 20 °C for autoinduction protein expression. Cells were then pelleted by centrifugation at 4,000 rpm for 20 minutes at 4 °C, and resuspended on ice in 30 mL lysis buffer (20 mM HEPES, 150 mM NaCl, 1 mM EDTA, 0.1 mM PMSF, 0.1 mg/mL lysozyme, pH 7.4). The resuspension was sonicated on ice for about 5 minutes to break the cells. After centrifugation at 13,000 rpm for 20 minutes at 4 °C, the supernatant was loaded onto ∼3 mL Ni-NTA column, and allowed to flow through by gravity. The column was washed with 50 mL wash buffer 1 (20 mM HEPES, 150 mM NaCl, pH 7.4) and wash buffer 2 (20 mM HEPES, 50 mM imidazole, 150 mM NaCl, pH 7.4). The his-tagged proteins were then eluted with 10 mL of elution buffer (20 mM HEPES, 150 mM NaCl, 200 mM imidazole, pH 7.4). The eluted proteins were further purified via a HW-55S size exclusion column in buffer (20 mM HEPES, 150 mM NaCl, pH 7.4) using an AKTA-purifier and fractions containing corresponding proteins were isolated. 20 mM TCEP was included in all buffers when preparing NS2B-D76C*/NS3-K117C.

To remove the his-tag from NS2B/NS3, NS2B/NS3 and TEV concentrations were determined by A280 using a NanoDrop 2000 (Thermo Scientific, Waltham, MA) with the theoretical molar absorption coefficient ε=52035 M^-1^⋅cm^-1^ and ε=39880 M^-1^⋅cm^-1^ respectively. The cleavage reaction was conducted by incubating TEV protease with NS2B/NS3 at a 1:10 molar ratio, in buffer (20 mM HEPES, 150 mM NaCl, pH 7.4) containing 10 mM DTT overnight at 37 °C. The samples (1mL) were incubated with Ni-NTA beads (0.1 mL) on a rotator for 10 minutes to remove the un-cleaved NS2B/NS3 and histagged TEV. Reaction efficiency and purity of his-tag cleaved products were analyzed on SDS-PAGE (Supplemental Figure S9). Samples were stored at -80 °C for later use. For NS2B-D76C*/NS3-K117C, 2 mM DTT was added in the storage buffer.

### NS2B-C76C*/NS3-K117C crosslinking

Immediately as NS2B/NS3 proteins eluted out from the HW-55S size exclusion column, 200 μM BMOE from 10 mM freshly prepared stock in DMSO was added to each elution fraction. Cross-linking reactions were incubated at room temperature on a tube rotator for 30 minutes before the reaction was quenched by the addition of 10 mM DTT. The crosslinking reaction samples were analyzed via band shifts on SDS-PAGE, where crosslinked proteins migrated noticeably faster than non-crosslinked (Supplemental Figure 10). Buffers of the crosslinked samples were then exchanged to buffer (20 mM HEPES, 150 mM NaCl, pH 7.4) by a size exclusion column HW-55S for storage at -80 °C.

### NS2B/NS3 activity assay

The NS2B/NS3 protease activity was determined by monitoring the fluorescence increase caused by cleaving a Pyr-RTKR-AMC peptide substrate. The assay was conducted in buffer (150 mM NaCl, 20 mM HEPES, pH 7.4), using 150 nM of NS2B/NS3 and 50 μM of Pyr-RTKR-AMC. Cleavage reactions were conducted in triplicate and monitored using 360/460 nm excitation/emission using a Synergy 2 Multi-Mode Microplate Reader (BioTek, Winooski, VT).

### Preparation of ClyA oligomers

ClyA protein concentrations were assessed with A280 using the theoretically determined molar absorption coefficient ε= 30370 M-1⋅cm-1. ClyA oligomers were prepared by incubating ClyA monomer at 25 μM with 1% DDM. For *St*ClyA-AS, oligomer samples were directly used in electrophysiology. For *Ec*ClyA-WT, a further gel extraction step was necessary to achieve nanopores of sufficient quality for electrophysiology. *E. coli* ClyA oligomers were loaded onto BioRad 4-20% Blue-Native gels. Gels were initially run for 60 minutes at 300 mV with an anode buffer of 50 mM bis-tris pH 7 and a dark cathode buffer of 15 mM bis-tris, 50 mM tricine, 0.02% G250 pH 7. The dark cathode buffer was then replaced with light cathode buffer of 15 mM bis-tris, 50 mM tricine pH 7, and run for another 120 minutes at 180 mV. The lowest oligomer band, corresponding to the dodecameric Type 1 ClyA pores used in our study, was excised and the proteins were extracted by 200-400 μL of buffer 150 mM NaCl, 20 mM HEPES, pH 7.4 with 0.1% DDM.

### Single channel current recording

Single-channel recording experiments were performed in an apparatus containing two chambers separated by a 25 μm thick Teflon film at 23 °C. An aperture of approximately 70-90 μm diameter was generated near the center of the film with an electric spark. The aperture was pretreated with a hexadecane in pentane (10% v/v) solution. After allowing the pentane to evaporate, each chamber was filled with 400 µL buffer (20 mM HEPES, 150mM NaCl, pH 7.4). An Ag/AgCl electrode was immersed in each chamber with the cis chamber grounded. DPhPC in pentane (10 mg/ml) was deposited on the surface of the buffer and monolayers were formed after the pentane evaporated. The lipid bilayer was formed by raising the liquid level up and down across the aperture. Capacitance was monitored to observe the formation of a functional membrane, which was 60-90 pF, depending on aperture size. Protein nanopore samples were added to the cis chamber and a voltage protocol was applied to facilitate the insertion. For the trapping experiment, *St*ClyA-AS of ∼-161 pA, *St*ClyA-E57A of ∼-146 pA, *St*ClyA-E57K of ∼-127 pA and *Ec*ClyA-WT of ∼-177 pA at -120 mV were used. NS2B/NS3 protease analytes were added to the cis chamber. After mixing, voltage potentials as specified in each experiment were applied across the bilayer and current signals were collected by a patch clamp amplifier (Axopatch 200B, Molecular Devices) and digitized by an analog-to-digital converter (Digidata 1440A, Molecular Devices) at a sampling rate of 50 kHz after processing with a 4-pole lowpass Bessel filter at 5 kHz. Data was recorded by Clampex 10.7 (Molecular Devices). Single-channel current recording traces were analyzed by Clampfit 11.2. Single-channel search of raw traces was used to collect the dwell times and the interevent time of captured events and a subsequent single-exponential fitting of a cumulative histogram of these data determined the τ_off_ and τ_on._ Example fittings are shown in Supplemental Figure S11. Raw traces were filtered by a low-pass100 Hz Gaussian filter and determine the dwell times of S_0_ and S_1_ states within captured events, after which double- or single-exponential fittings of cumulative histograms of the data determined the τ_off_ of each state. Example fittings carried out in Origin are shown in Supplemental Figure S12.

### Computer modeling of the WNV protease/ClyA system

Considering the large size and complexity of the protease and nanopore system, we applied the hybrid resolution (HyRes) coarse-grained model ^56, 57^ to allow for sufficient sampling of potential dynamic and non-specific interactions between the NS2B/NS3 protease and the ClyA pore. HyRes has an atomistic backbone and up to 5 bead resolution for sidechain representations, which is parameterized to reproduce the geometry and effective interactions of each amino acid. The model has been successfully applied to study dynamic interactions between disordered proteins and/or folded domains, including spontaneous phase separations ^58, 59^. Therefore, the HyRes model is expected to be appropriate to study the dynamic interactions of WNV protease in ClyA nanopore.

The initial HyRes structures of the ClyA nanopore and WNV NS2B/NS3 protease were derived from atomistic PDB structures (using at2hyres script at https://github.com/mdlab-um/HyRes_GPU). PDB:2WCD was used for *E. coli* ClyA in the 12-mer form. Unstructured C- and N-terminal tails (11 and 7 residues, respectively) were not included in HyRes modeling, because they are far away from the pore lumen region. During the simulations, all Cα atoms on ClyA have been harmonically restrained to their original positions with a force of 1.0 kcal/mol to maintain the structure. All other ClyA constructs were derived from the WT *E. coli* ClyA model through computational mutagenesis. For the WNV protease, PDB:2IJO was used for the closed state. The G_4_SG_4_ linker was added to connect NS2B and NS3 protease to be consistent with the experimental conditions. Distance-based restrains were applied to the structured region of the protease with a force constant of 0.5 kcal/mol to maintain the structure, except that the restrains were removed for the NS2B CTD (residue 48 to 59) in the open state. These restrains were able to maintain the protease structures and capture the difference in closed and open states (see Supplementary Figure S13).

All simulations were carried out using CHARMM ^66, 67^ with constant volume and temperature (NVT) conditions. The SHAKE algorithm ^68^ was used to constrain all bonds involving hydrogen atoms to allow for a 2-fs integration step with Langevin thermostat. In all simulations, the short-range non-bonded interactions were smoothly truncated at 1.8 nm. The Debye-Hückel method was used to treat long-range electrostatic interactions with a salt concentration of 0.15 M.

### Pulling and standard MD simulations of the protease/ClyA interactions

We first executed the steered MD simulations to explore the potential interactions between pore and protease. The protease was initially placed 4.5 nm away from the nanopore in 10 different rotations, with the center of mass facing the center of the ClyA pore lumen (Figure S3). This setup aimed to avoid the simulation bias from initial configuration and allow for diverse orientation of protease when entering the pore, since the cis entrance of ClyA is larger than the longest dimension of protease. The pulling force was added to the Cα atom of residue H122 on protease NS3, which is close to the center of mass, with a moderate 60 pN constant directing to the nanopore center (cis-trans direction, Figure S3). All simulations were carried out at 300 K and lasted for 50 ns.

We then performed the standard MD simulations to specifically explore the protease dynamics in the center of *Ec*ClyA-WT, *St*ClyA-AS, *St*ClyA-E57A, and *St*ClyA-E57K pore variants, respectively. Particularly, only residues 36-86, 123-176 and 208-256 on ClyA have been kept, which are in the reference to the “mid-region” of ClyA lumen. Protease were placed in the center of ClyA with the center of mass in line with the pore center. Six different orientations were used as initial conformations, where the NS2B CTD faces up, down, right, left, front, and back to the ClyA pore, to enable more sufficient sampling (Supplementary Figure S14). Each orientation was running with two replicas. All simulations were performed at 400 K, where a higher temperature was used to accelerate the simulation and lower the probability of some unfavorable protease confinements in the pore. All simulations last for 200 ns.

### Analysis of MD-generated conformational ensembles

During the pulling simulations, the z-depth measured the center of mass of NS2B/NS3 protease, where the origin (z = 0) was set at the center of the membrane (Figure S3). The z-direction aligns with the trans-cis direction. VMD is used for protein visualization^69^.

In the standard MD simulations, the first 100 ns was excluded for all 200-ns coarse-grained simulation trajectories in the subsequent analysis. All analyses were performed with a combination of CHARMM, in-house scripts, and the python codes with MDTraj package^70^. The principal component analysis (PCA) was first performed using python SciKit-learn package^71^, to evaluate the sampling convergence as well as to visualize the simulated ensembles. For this, snapshots were taken every 100 ps from the entire 100 ns trajectories to collect all sampled protease conformations. Particularly, the symmetric conformation of protease along z direction has been aligned to one single direction to avoid over-counting. The ensembles generated from all simulations (6 orientations of protease in four ClyA pore with two replica each orientation, 24 trajectories in total) were combined together and then using the coordinates of Cα atoms except for the NS2B CTD for the PCA analysis. This analyses captured how protease interact with ClyA, including the binding position and adopted orientations. The free energy surfaces shown were derived directly from the 2D probability distributions along the first two principal components (PCs).

To estimate the solvated ion cross-section area and residual current (*I*_res_%) of protease in the pore, the in-house CHARMM scripts were used to count the whole area of ClyA lumen, and the rest area being occupied by protease. The 3-dimensional pore lumen was first divided into multiple layers with each layer’s height up to 0.1 Å. The counting of the surface area in each layer was done by subtracting the area occupied by protein beads from a square. For the non-constriction region in the pore lumen, the width of the square is 50 Å (the longest lumen diameter is ∼ 55 Å). For the constriction region, the width of the square is 40 Å (the shortest lumen diameter is ∼ 35 Å). The water probe is set to 1.4 Å. As for the residual current calculation, *I*_res_% is inversely proportional to the resistance, while the resistance can be calculated from the cross-section by integrating the cross-section through the whole pore,

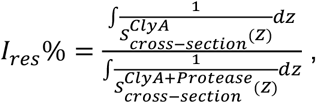

where 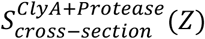 stands for the cross section area with protease in the pore and 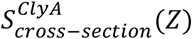 stands for the cross section of ClyA whole lumen without protease.

