## Supplemental figures for "Tuning single-molecule ClyA nanopore tweezers for real-time tracking of the conformational dynamics of West Nile viral NS2B/NS3 protease"

**CONTENTS**

**Figure S1:** Sequence alignment of EcWT and StAS ClyA……………………………………..……..2

**Figure S2:** Extended traces of WT protease in StAS and EcWT ClyA………………….................3

**Figure S3:** Configuration of steered MD simulations………………………………………….......4

**Figure S4:** Trajectories and interactions of steered MD simulations ………………………………5

**Figure S5**: Cartoon representation of the simulated mid-trap states in ClyA pores ……………...6

**Figure S6:** Activity assay results for protease constructs ……………………………...................7

**Figure S7:** Extended traces of protease constructs in StE57A …………………………….............8

**Figure S8:** Example trace of a rare long dwell time event for rd-C76*/C117……………………..…9

**Figure S9:** SDS-PAGE gels for protease constructs ……………………………….………………10

**Figure S10:** Crosslinking SDS-PAGE gel for xc-C76*/C117Purification ………………………...11

**Figure S11:** Sample analysis fittings to derive $\tau_{off}$ and $\tau_{on}$……………………..……………………12

```
E. coli ClyA      MTEIVADKTVEVVKNAIETADGALDLYNKYLDQVIPWQTFDETIKELSRFKQEYSQAASV 60
S. typhi ClyA-AS  MTGIFAEQTVEVVKSAIETADGALDLYNKYLDQVIPWKTFDETIKELSRFKQEYSQEASV 60
                  ** *.*::******.**********************:****************** ***

E. coli ClyA      LVGDIKTLLMDSQDKYFEATQTVYEWCGVATQLLAAYILLFDEYNEKKASAQKDILIKVL 120
S. typhi ClyA-AS  LVGDIKVLLMDSQDKYFEATQTVYEWAGVVTQLLSAYIQLFDGYNEKKASAQKDILIRIL 120
                  ******.*******************.**.****:*** *** **************::*

E. coli ClyA      DDGITKLNEAQKSLLVSSQSFNNASGKLLALDSQLTNDFSEKSSYFQSQVDKIRKEAYAG 180
S. typhi ClyA-AS  DDGVKKLNEAQKSLLTSSQSFNNASGKLLALDSQLTNDFSEKSSYYQSQVDRIRKEAYAG 180
                  ***:.**********.*****************************:*****:********

E. coli ClyA      AAAGVVAGPFGLIISYSIAAGVVEGKLIPELKNKLKSVQNFFTTLSNTVKQANKDIDAAK 240
S. typhi ClyA-AS  AAAGIVAGPFGLIISYSIAAGVVEGKLIPELNNRLKTVQNFFTSLSATVKQANKDIDAAK 240
                  ****:**************************:*:**:******:** *************

E. coli ClyA      LKLTTEIAAIGEIKTETETTRFYVDYDDLMLSLLKEAAKKMINTCNEYQKRHGKKTLFEV 300
S. typhi ClyA-AS  LKLATEIAAIGEIKTETETTRFYVDYDDLMLSLLKGAAKKMINTSNEYQQRHGRKTLFEV 300
                  ***:******************************* ********.****:***:******

E. coli ClyA      PEVHHHHHH--- 309
S. typhi ClyA-AS  PDVGSSYHHHHH 312
                  *:*   :**
```

**Figure S1.** Sequence alignment of *E. coli* and *S. typhi* ClyA-AS with the key luminal residue differences highlighted. The sequences have a percent identity of 88.4%.

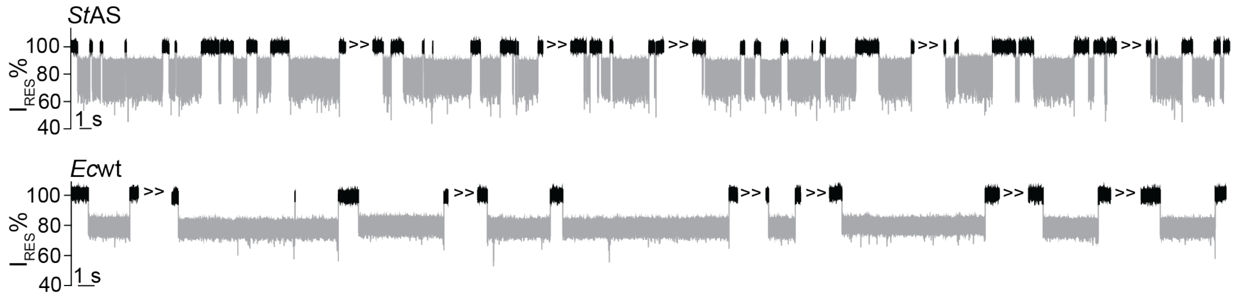

**Figure S2.** Extended traces showing WT protease trapping by *S. typhi* ClyA-AS and *E. coli* ClyA. The traces were recorded at -120 mV in 150 mM NaCl, 20 mM HEPES pH 7.4 buffer at a sampling rate of 50 kHz and a lowpass filter rate of 5 kHz.

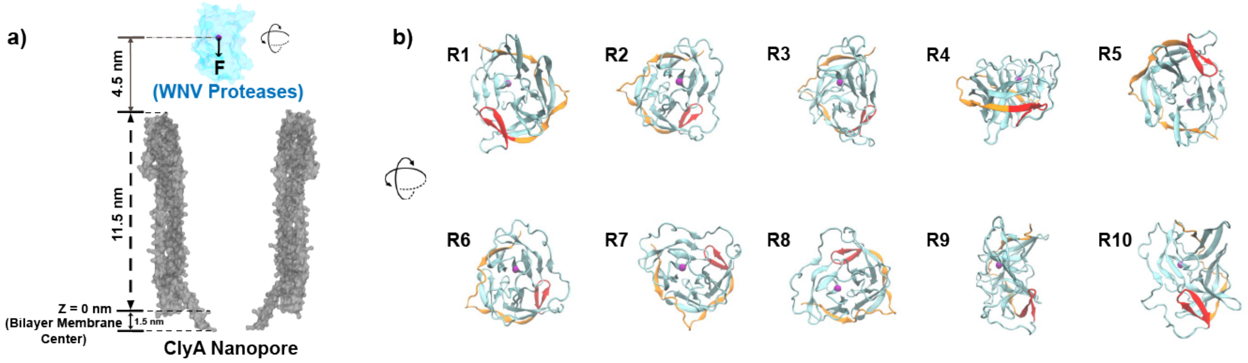

**Figure S3.** Configuration of steered MD simulations of the protease-ClyA interactions. **a**) Illustration of initial configuration of steered MD simulations, where the protease was placed ~4.5 nm away from the extracellular entrance of the ClyA nanopore. A constant pulling force of 60 pN was at applied to the center of mass. **b**) The 10 different initial orientations of protease for pulling simulations. The protease is drawn in cartoon style, with NS3 highlighted in cyan, NS2B in orange, and the NS2B C-terminal hairpin in red.

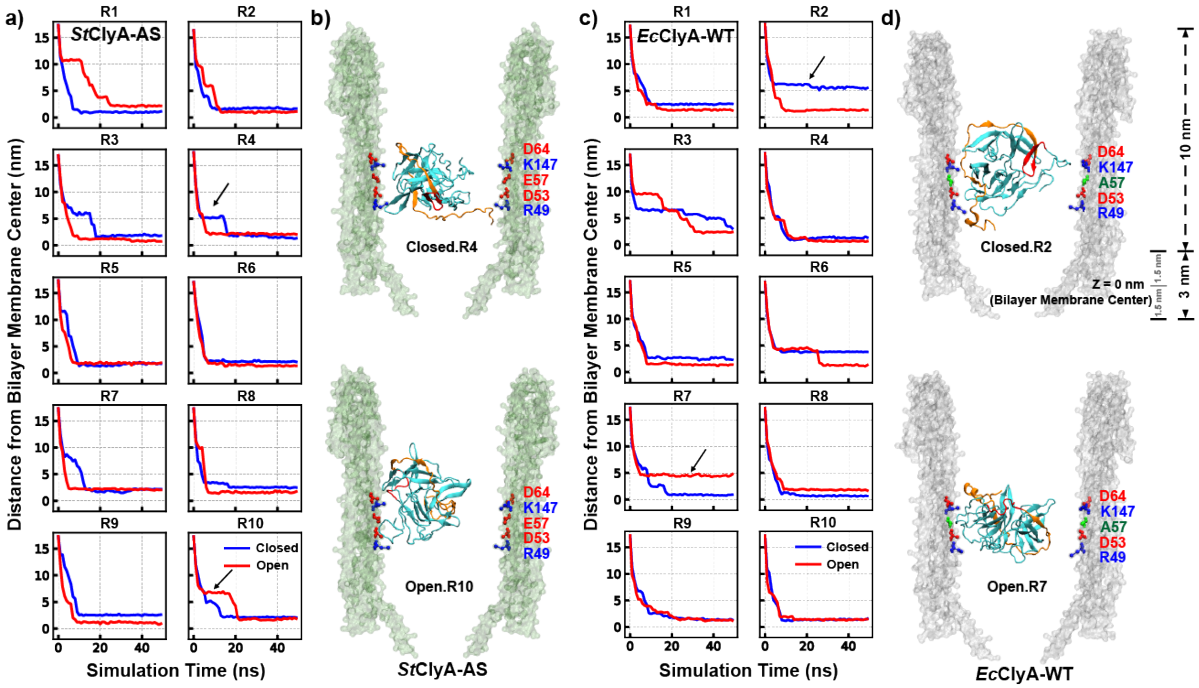

**Figure S4.** Pulling simulations of the WNV protease in ClyA nanopores. a) The center of mass of the protease as a function of simulation time for *S. typhi* and c) *E. coli* ClyA nanopores, initiated from the 10 different orientations shown in Figure S3. The blue and red traces correspond to the protease in the open and closed conformations, respectively. Snapshots of representative mid-trap states are shown in b) and d), with the corresponding time points marked with arrows in panels a) and c).

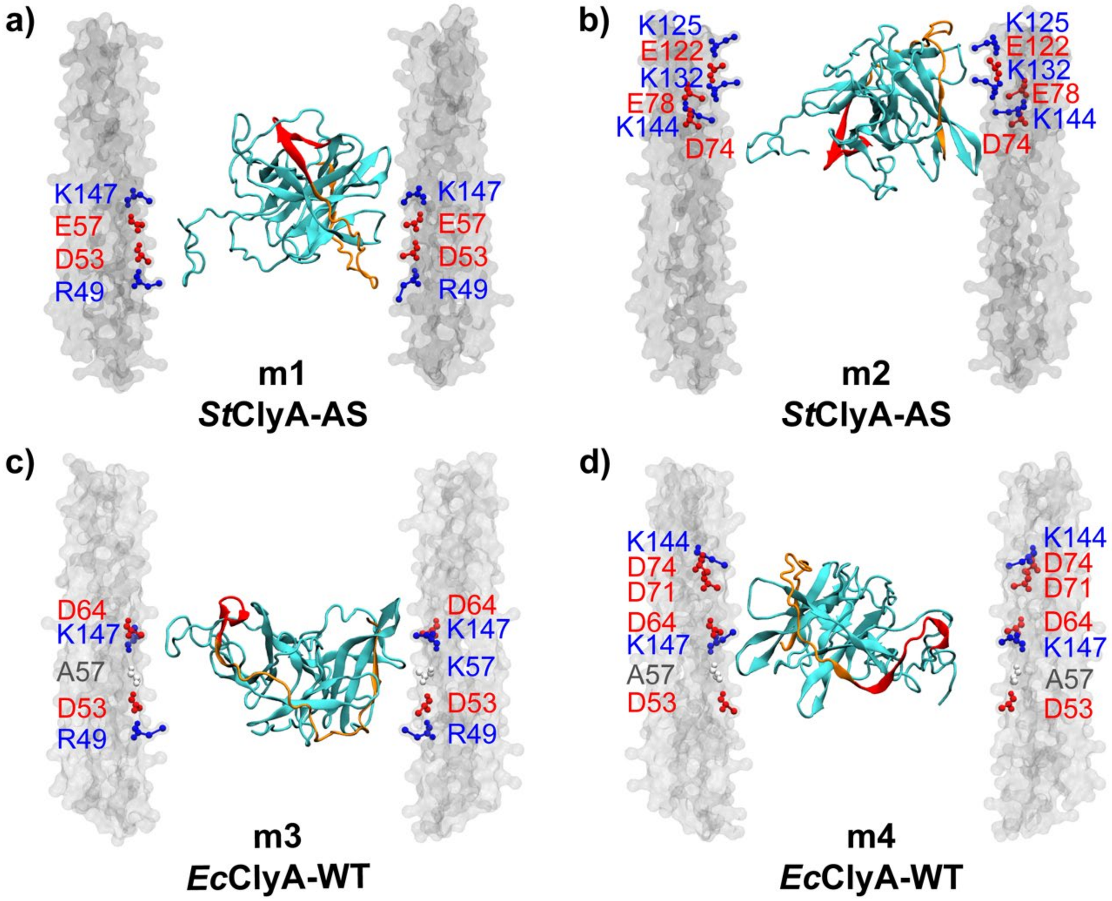

**Figure S5**: Cartoon representation of the mid-trap states in ClyA pores. The protease is drawn following the same style as Figure S3b. Key binding residues on the nanopore are labeled as in Figure 2c.

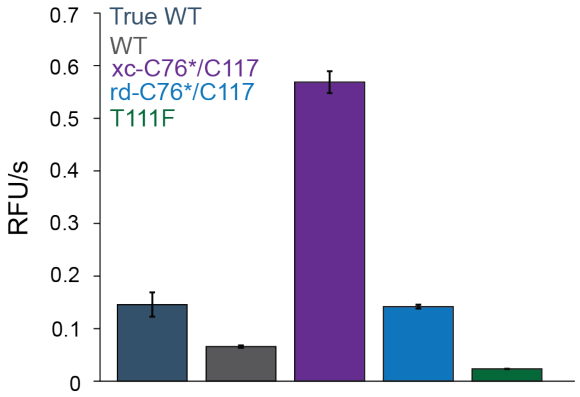

**Figure S6.** Activity assay results for the True WT, WT, xc-C76*/C117, rd-C76*/C117, and T111F protease constructs with the peptide sbustrate Pyr-RTKR-AMC. WT refers to the cysteineless construct with a C78A mutation, while True WT does not have this mutation. The rates in RFU per second are plotted for the condition of 150 nM enzyme and 50 μM substrate.

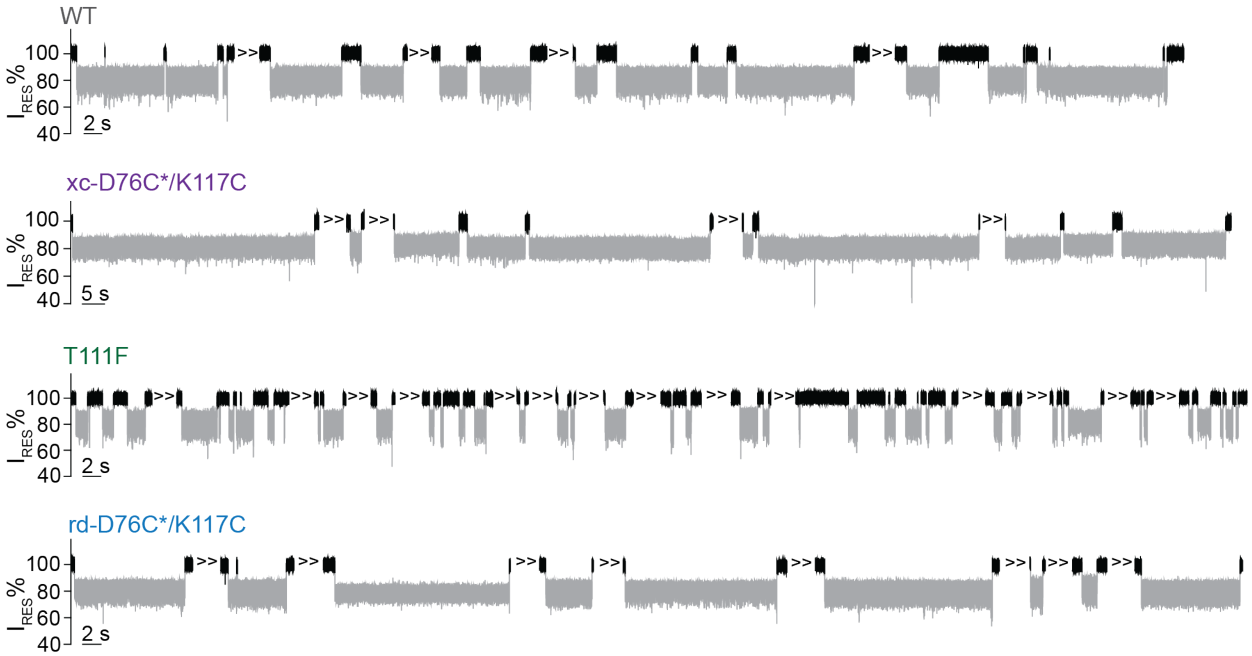

**Figure S7.** Extended traces showing WT, xc-C76*/C117, T111F, and rd-C76*/C117 variants of the WNV NS2B/NS3 protease trapping by *St*ClyA-E57A. The traces were recorded at -120 mV in 150 mM NaCl, 20 mM HEPES pH 7.4 buffer at a sampling rate of 50 kHz and a lowpass filter rate of 5 kHz.

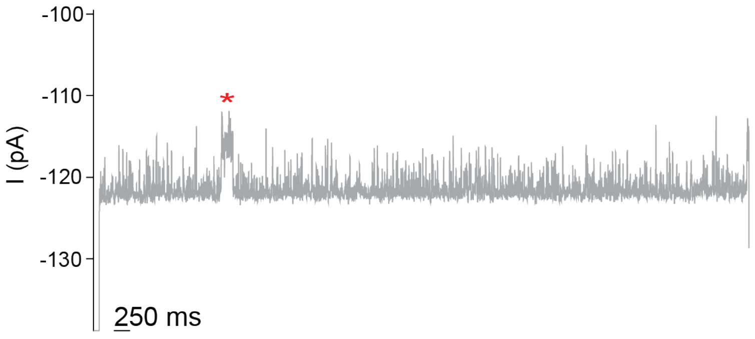

**Figure S8.** A current trace of rd-C76*/C117 showing a rare long dwell time event. The current recording was performed at -120 mV in 150 mM NaCl, 20 mM HEPES pH 7.4 buffer at a sampling rate of 50 kHz and a lowpass filter rate of 5 kHz. The trace was filtered with a 100 Hz low-pass Gaussian filter.

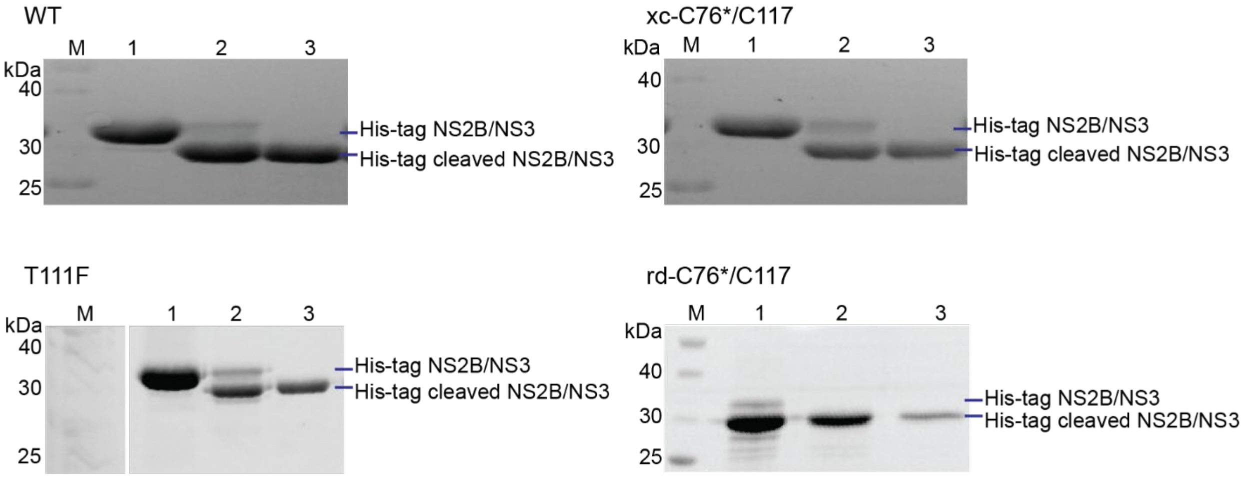

**Figure S9.** SDS-PAGE of the WT, xc-C76*/C117, T111F, and rd-C76*/C117 protease constructs following purification. For the gels of WT, xc-C76*/C117, and T111F lane 1 is unreacted NS2B/NS3, lane 2 is following TEV digestion, and lane 3 is the final purified NS2B/NS3 product. For the rd-C76*/C117 gel, lane 1 is TEV digested NS2B/NS3, lane 2 is following NiNTA pulldown, and lane 3 is purified NS2B/NS3 product.The upper band corresponds to uncleaved NS2B/NS3 and the lower band corresponds to successfully cleaved NS2B/NS3.

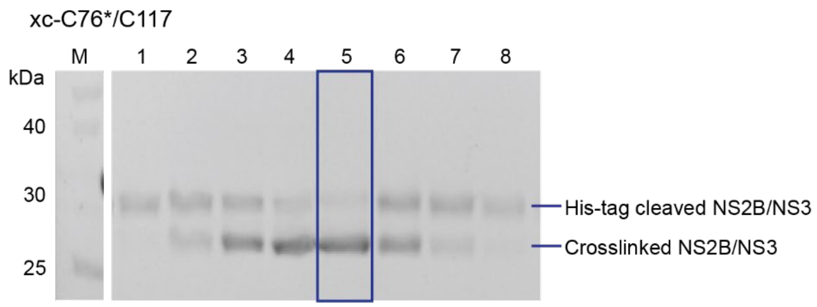

**Figure S10.** SDS-PAGE of the crosslinking reaction of NS2B D76C/NS3 K117C, C78A with BMOE to fully generate xc-C76*/C117. Fractions 1-8 are shown, and a crosslinking efficiency of 90% was achieved for fraction 5. The upper band corresponds to cleaved NS2B/NS3 and the lower band corresponds to successfully crosslinked NS2B/NS3.

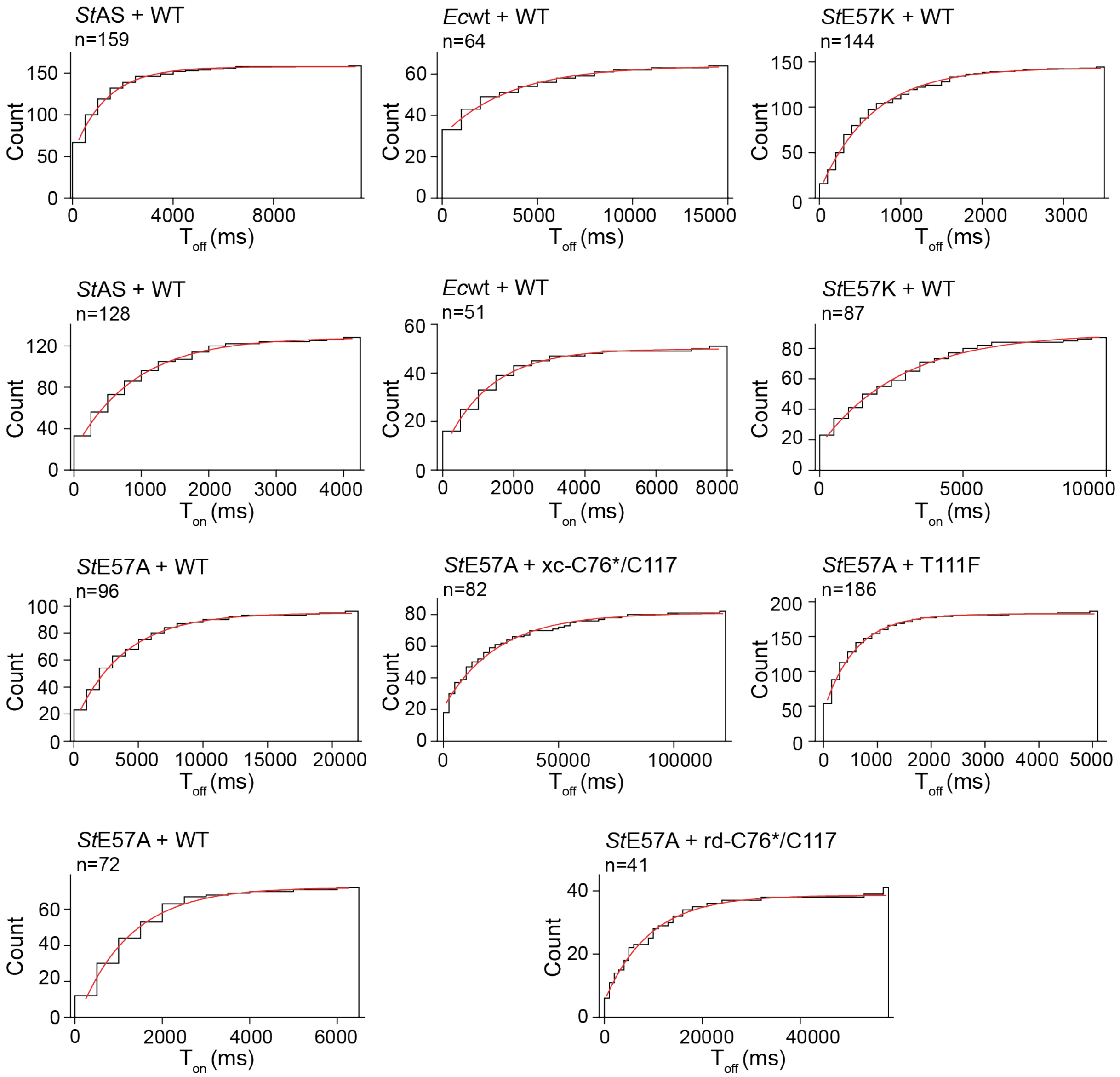

**Figure S11.** Sample fittings from Clampfit 11.2 of the collected data used to determine $\tau_{off}$ and $\tau_{on}$ at -120 mV. The number of events (n) fit is indicated for each.

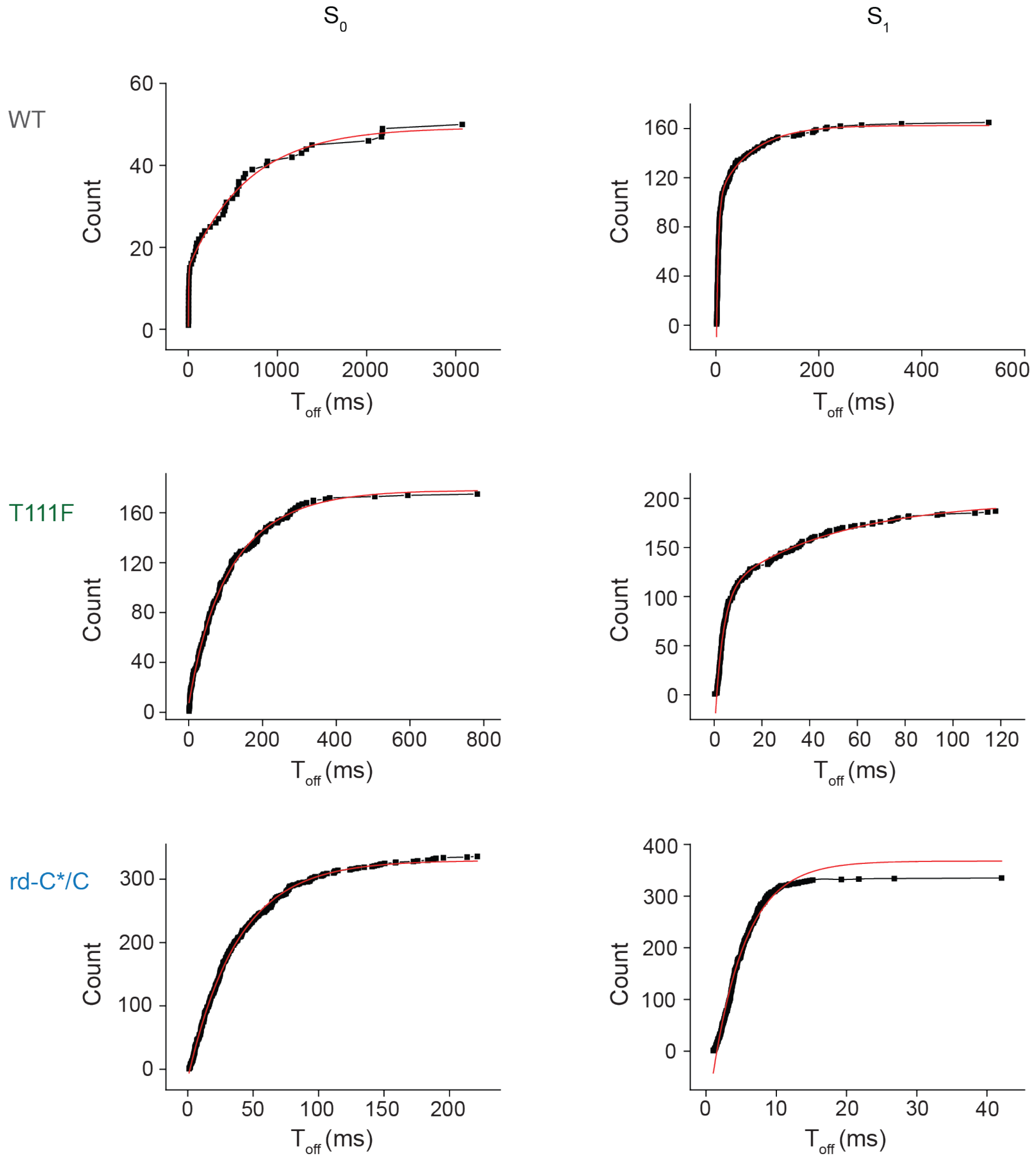

**Figure S12.** Dwell time analysis of the $S_0$ and $S_1$ states. Raw traces were filtered by a 100 Hz Gaussian filter and analyzed by Clampfit11.2 to select the $S_0$ and $S_1$ events via the Single Channel Search function. Cumulative histograms of the dwell times of events were fitted with double-exponential or single exponential functions in Origin.

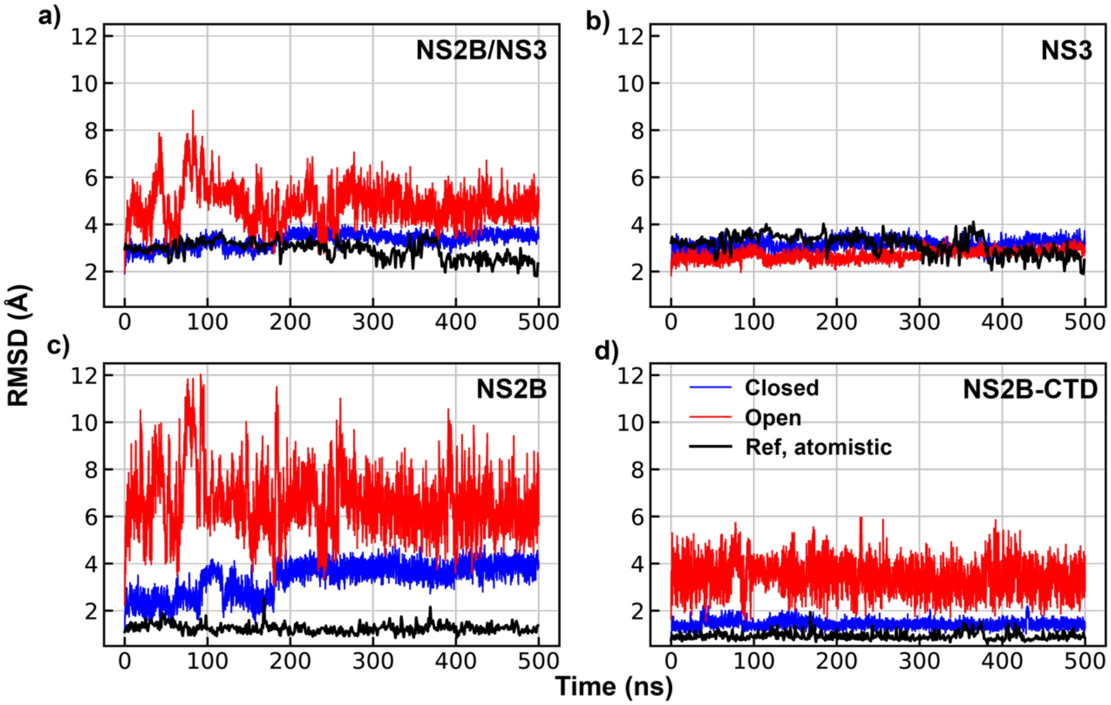

**Figure S13.** Stability of the WNV NSB2/NS3 protease in HyRes with structural restraints. The atomistic simulation was performed using the CHARMM36m explicit solvent force field. The open state is much more flexible in HyRes simulations mainly due to the NS2B C-terminus, which can rapidly unfold in HyRes but not in atomistic simulations due to the simulation timescale.

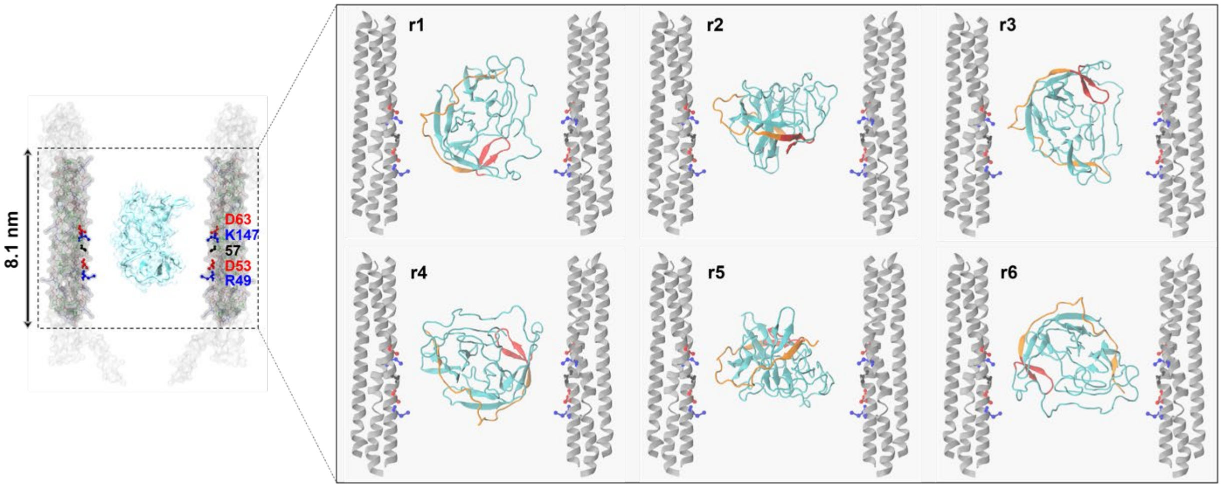

**Figure S14.** Standard MD simulations of protease/ClyA interactions. Only the highlighted region of ClyA was included, while the transparent regions were deleted to reduce the computational cost. The protease was initially placed in the middle of the pore with six different orientations (r1 to r6). The protease was represented using the same style as in Figure S4.

**Table S1.** Primers for ClyA and NS2B/NS3 protease variants cloning.

| Mutants | Primer Sequences |
|---|---|
| *St*ClyA-E57A | AGTCAAGCAGCGTCCGTCCTAGTG |
| | GGACGCTGCTTGACTGTATTCCTG |
| *St*ClyA-E57K | AGTCAAAAAGCGTCCGTCCTAGTG |
| | GGACGCTTTTTGACTGTATTCCTG |
| NS3-C78A | CGACTTGCCTACGGAGGACCCTGG |
| | TCCGTAGGCAAGTCGATCCTCTTTG |
| NS2B-D76C | AGAGTTTGTGTGCGGCTTGATGATG |
| | CCGCACACAAACTCTCTCGCTCGAG |
| NS3-K117C | GTGTTCTGTACACCTGAAGGAGAAATTG |
| | AGGTGTACAGAACACCCCTGGTTTC |
| NS3-T111F | GTCCAGTTCAAACCAGGGGTGTTC |
| | TGGTTTGAACTGGACGTTTTTAAC |

**Table S2.** Sequences of NS2B/NS3 protease constructs.

| Construct | Sequence |
|---|---|
| True WT | MGSSHHHHHHSSENLYFQSTDMWIERTADISWESDAEITGSSERVDVRLDDDGNFQL<br>MNDPGAQYTGGGGSGGGGGGVLWDTPSPKEYKKGDTTTGVYRIMTRGLLGSYQAG<br>AGVMVEGVFHTLWHTTKGAALMSGEGRLDPYWGSVKEDRLCYGGPWKLQHKWNG<br>QDEVQMIVVEPGKNVKNVQTKPGVFKTPEGEIGAVTLDFPTGTSGSPIVNKNGDVIGL<br>YGNGVIMPNGSYISAIVQGERMDEPIPAGFEPEMLRK |
| WT | MGSSHHHHHHSSENLYFQSTDMWIERTADISWESDAEITGSSERVDVRLDDDGNFQL<br>MNDPGAQYTGGGGSGGGGGGVLWDTPSPKEYKKGDTTTGVYRIMTRGLLGSYQAG<br>AGVMVEGVFHTLWHTTKGAALMSGEGRLDPYWGSVKEDRLAYGGPWKLQHKWNG<br>QDEVQMIVVEPGKNVKNVQTKPGVFKTPEGEIGAVTLDFPTGTSGSPIVNKNGDVIGL<br>YGNGVIMPNGSYISAIVQGERMDEPIPAGFEPEMLRK |
| xc- and rd-C76/C117 | MGSSHHHHHHSSENLYFQSTDMWIERTADISWESDAEITGSSERVCVRLDDDGNFQL<br>MNDPGAQYTGGGGSGGGGGGVLWDTPSPKEYKKGDTTTGVYRIMTRGLLGSYQAG<br>AGVMVEGVFHTLWHTTKGAALMSGEGRLDPYWGSVKEDRLAYGGPWKLQHKWNG<br>QDEVQMIVVEPGKNVKNVQTKPGVFCTPEGEIGAVTLDFPTGTSGSPIVNKNGDVIGL<br>YGNGVIMPNGSYISAIVQGERMDEPIPAGFEPEMLRK |
| T111F | MGSSHHHHHHSSENLYFQSTDMWIERTADISWESDAEITGSSERVDVRLDDDGNFQL<br>MNDPGAQYTGGGGSGGGGGGVLWDTPSPKEYKKGDTTTGVYRIMTRGLLGSYQAG<br>AGVMVEGVFHTLWHTTKGAALMSGEGRLDPYWGSVKEDRLCYGGPWKLQHKWNG<br>QDEVQMIVVEPGKNVKNVQFKPGVFKTPEGEIGAVTLDFPTGTSGSPIVNKNGDVIGL<br>YGNGVIMPNGSYISAIVQGERMDEPIPAGFEPEMLRK |

*Mutations were highlighted in red.
